# Targeting the APRIL-BAFF axis in IgA nephropathy: preclinical insights from influenza challenge and single-cell spatial profiling

**DOI:** 10.64898/2026.05.01.722321

**Authors:** Kirk J Rowley, Vinay Singh, Anna Roberts, Katherine A Halley, Joshua Brand, Jacob Vinay Konakondla, Daved Fared, Michelle Lu, Brett Hurst, Wayne W Hancock, Gregory J Babcock, Luke N Robinson

## Abstract

**Background:** APRIL and BAFF are TNF superfamily cytokines that regulate B-cell development, survival, and antibody production, and are emerging therapeutic targets for IgA nephropathy (IgAN). Selective APRIL and dual APRIL/BAFF inhibitors both reduce IgA and proteinuria in IgAN clinical trials, but whether their broader immunological consequences differ has not been systematically characterized.

**Methods:** We compared selective APRIL and dual APRIL/BAFF inhibition using influenza vaccination and lethal challenge, KLH immunization, serological profiling, and flow cytometry in mice, alongside human B-cell survival assays in vitro. Single-cell CITE-seq and in situ spatial transcriptomics were applied to characterize molecular and tissue-level changes in the spleen.

**Results:** Both modes of inhibition reduced serum IgA by ≥60% in mice. However, dual APRIL/BAFF inhibition nearly abolished vaccine-mediated protection against lethal influenza challenge (10% versus 70% survival in controls; p < 0.01), whereas selective APRIL inhibition had limited impact on protective immunity. This functional divergence was underpinned by broad cellular disruption under dual blockade, including >80% depletion of splenic B cells, loss of T follicular helper cells, and impaired antigen-specific IgM and IgG responses. Selective APRIL inhibition left these populations and responses largely intact. Consistent with these findings, human B-cell survival in vitro was dependent on BAFF, not APRIL. Single-cell and spatial transcriptomics revealed that dual blockade collapsed follicular architecture, eliminated germinal centers, and disrupted chemokine organization, whereas these structures remained intact under selective APRIL inhibition. At the molecular level, dual blockade, but not selective APRIL inhibition, downregulated NF-κB survival signaling and antigen presentation programs and shifted surviving germinal center B cells toward a pro-apoptotic state.

**Conclusions:** Selective APRIL and dual APRIL/BAFF inhibition both reduce IgA, the pathologically relevant isotype in IgAN, but only dual blockade disrupts B-cell maturation, germinal center function, tissue architecture, and protective immunity. These findings inform the benefit-risk assessment of chronic B cell-targeting therapies in IgAN.

## INTRODUCTION

Immunoglobulin A (IgA) nephropathy (IgAN) is the most common primary glomerulonephritis worldwide with approximately 200,000 people afflicted in the United States^1, 2^ and a global incidence of about 2.5 per 100,000 people per year^3^. The disease ultimately leads to kidney failure in a large percentage of patients^2^. The pathogenesis of IgAN is explained by a “four-hit” process initiated by elevated circulating levels of pathogenic molecular variants of polymeric IgA1 containing an aberrantly O-glycosylated hinge region (Gd-IgA1), followed by synthesis of autoantibodies against Gd-IgA1, immune complex formation and deposition of these complexes in the mesangium^4, 5^. This culminates in infiltration of inflammatory cells and complement activation, resulting in hematuria, proteinuria, glomerular sclerosis, and ultimately, progression to kidney failure.

Therapy for IgAN historically has been limited to renin-angiotensin system (RAS) blockade with the use of angiotensin converting enzyme (ACE) inhibitors, angiotensin receptor blockers (ARBs) and gluticoticoids^6^. Therapeutic strategies in IgAN are rapidly evolving, with several approved products since 2021, and multiple emerging treatments in clinical development targeting different aspects of IgAN pathogenesis. Approved molecules include targeted-release corticosteroids, endothelin receptor antagonist/blocker, complement inhibitors and APRIL inhibitor. Although these agents reduce proteinuria and slow eGFR decline, they do not fully address the underlying mechanisms driving disease progression, and many patients remain at risk for progressive kidney function loss^7^.

The B cell lineage plays an important role in the pathogenesis of several immune-mediated kidney diseases, including IgAN^8^, through overlapping mechanisms including autoreactive antibody-secreting cell development and propagation, antigen presentation, cytokine secretion and formation of autoantibodies and immune complex deposition. Multiple therapeutic targets along the B-cell development pathway can be modulated by different strategies. The cytokines APRIL (*TNFSF13*) and BAFF (*TNFSF13B*) are members of the TNF superfamily and are emerging therapeutic targets for modulating B-cell activity in kidney disease^9^. APRIL and BAFF are essential cytokines for the development and survival of cells within the B-cell lineage. Interestingly, APRIL mediates antibody class switching in mature B cells, leading to production of IgA,including pathogenic Gd-IgA1, and prolonging plasma cell survival^10^. BAFF has a broader impact on the B-cell compartment, supporting the survival of transitional and mature B cells, maintaining B-cell homeostasis, and promoting maturation of peripheral B cells^11^. APRIL and BAFF both interact with B cell maturation antigen (BCMA) and transmembrane activator and CAML interactor (TACI) receptors with varying affinity; however, the primary receptor for BAFF is the BAFF receptor (BAFF-R)^12–14^. BAFF-R is mainly expressed on peripheral immature and mature B cells, whereas TACI and BCMA are expressed on mature memory B cells and plasma cells^12, 15^. BAFF/BAFF-R interaction is an essential signaling event for the development and survival of peripheral B cells, as knockout of BAFF-R results in a near-complete loss of naïve B-cell development^16^. Knockout of TACI or BCMA has no impact on early B-cell development but leads to a reduction in antibody secreting cells and reduction of circulating immunoglobulin, most notably IgA in the case of TACI^17, 18^.

Converging lines of evidence from animal models, human genetics, and clinical studies point to APRIL as a central driver of IgAN pathogenesis, while the contribution of BAFF is less well established. In IgAN-prone gddY mice, anti-APRIL treatment reduces serum IgA, immune complex deposition in the kidney and proteinuria^19^, and APRIL-deficient mice display impaired IgA class switching. Clinically, elevated APRIL levels have been observed in patients with IgAN^20,21^ and these levels directly correlate with disease severity. Genome-wide association studies (GWAS) have further established TNFSF13 (APRIL) as a risk locus for IgAN susceptibility^22, 23^. In contrast, the evidence implicating BAFF in IgAN is less substantial. Although transgenic mice over-expressing BAFF develop elevated IgA, mesangial IgA deposition, hematuria, and proteinuria; however, GWAS analyses have not established a role for BAFF in IgAN susceptibility. A correlation between circulating BAFF levels and protenuria has been reported^24^, but treatment of IgAN patients with the anti-BAFF peptibody blisibimod did not result in a statistically or clinically meaningful resolution of disease^25^.

Therapies targeting these pathways in IgAN now include an approved anti-APRIL monoclonal antibody (sibeprenlimab) as well as dual APRIL/BAFF inhibitors in advanced clinical development^26–29^. Despite human genetic analysis and studies in animal models, the exact roles of APRIL and BAFF in IgA nephropathy have not been fully elucidated and requires further investigation, including the therapeutic potential of targeting APRIL or both APRIL and BAFF. Because APRIL and BAFF act at different stages of B-cell differentiation, their inhibition may have divergent impacts on disease. At the same time, the consequences for protective immunity may differ substantially, making this distinction important for evaluating the safety profile, including infection susceptibility, of emerging therapies. This is highlighted by multiple human clinical studies evaluating anti-BAFF or anti-APRIL/BAFF inhibitors demonstrating increased frequency of infection and lower vaccine response rates in some patients^30–32^.

Here we present data characterizing the differential effects of selective APRIL inhibition and dual APRIL/BAFF inhibition on immune B-cell populations, serological responses, and protective immunity in preclinical models. By clarifying how these pathways shape immune composition and function, this study seeks to provide mechanistic insights that may help inform therapeutic strategies for IgAN which directly treat the main drivers of disease while exhibiting the least impact on global immune responses.

## RESULTS

### Dual APRIL/BAFF but not APRIL-only inhibition impairs protective immunity to influenza

Given that APRIL and BAFF both play central roles in B-cell survival and antibody production, we first asked whether their therapeutic inhibition carries distinct functional consequences for protective immunity, before dissecting the underlying immunological mechanisms. To address this, we employed an influenza vaccination and live viral challenge model designed to test whether inhibitor-treated mice could mount and sustain a protective vaccine response (Figure 1A). Mice were dosed twice weekly for up to 13 weeks via intraperitoneal injection beginning at week 0 with one of four treatments: an APRIL inhibitor (antibody 4540), a dual APRIL/BAFF inhibitor (POV, a recombinant protein with the same amino acid sequence as povetacicept, produced internally; see Methods), an isotype control (MOTA), or vehicle control (PBS). At weeks 4 and 7, all mice, except the unvaccinated control group, received inactivated H1N1 Cal09 influenza virus vaccine. All mice were then challenged with live H1N1 Cal09 virus at week 10. As expected, unvaccinated mice uniformly succumbed to infection within 8 days of challenge, confirming the lethality of the challenge (Figure 1B). Both the PBS and isotype control groups achieved 70% survival, establishing a baseline for vaccine-mediated protection. Dual APRIL/BAFF inhibition nearly abolished this protection, reducing survival to 10% (p < 0.01 versus isotype control). Of note, the single surviving animal in the dual APRIL/BAFF group showed no body weight loss throughout the challenge, unlike the rapid decline observed in the remaining nine animals, raising the possibility that this mouse was not productively infected (Supplementary Figure S1). In contrast, APRIL-only inhibition resulted in only a modest reduction in survival that did not reach statistical significance relative to vehicle or isotype controls, suggesting that selective APRIL blockade largely preserves the ability to mount a functional vaccine response. Taken together, these results reveal a strong functional divergence between selective APRIL and dual APRIL/BAFF inhibition relating to their impact on protective immunity and prompted us to investigate the immunological basis for this difference.

**Figure 1.**
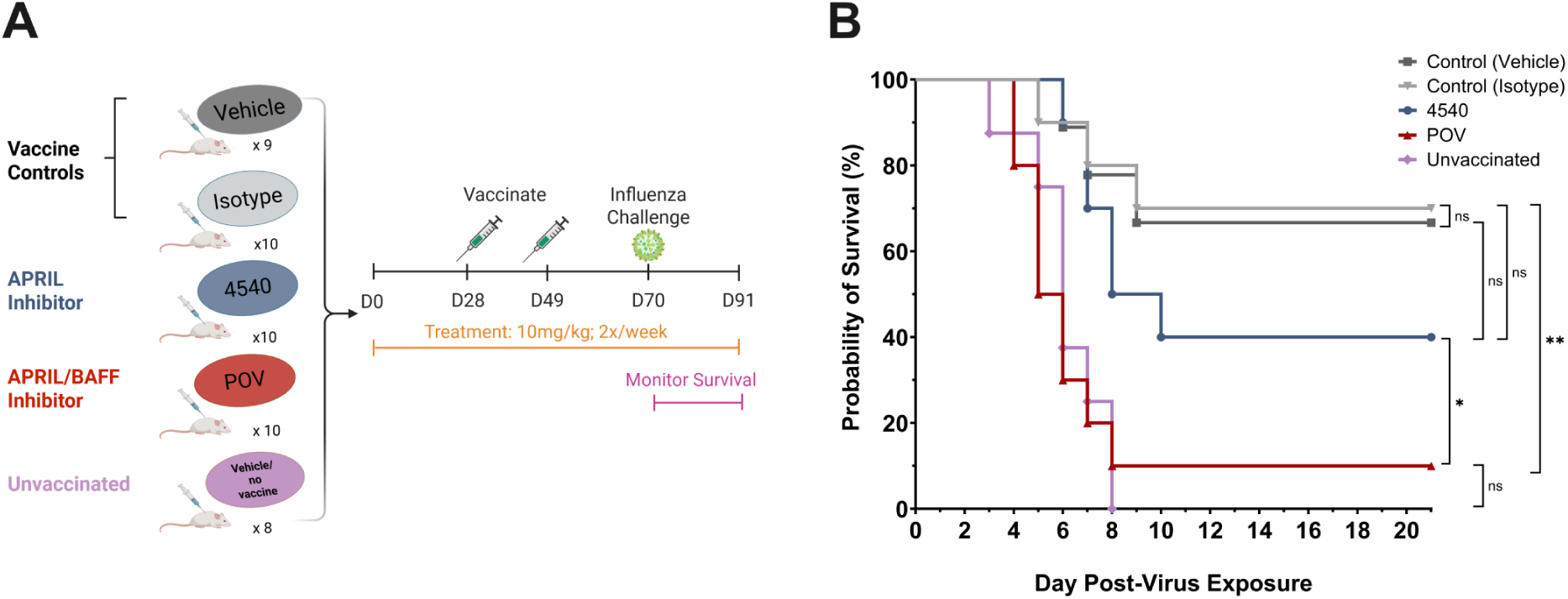
Differing effect of dual APRIL/BAFF inhibition versus APRIL-only inhibition on vaccine-mediated protection in a mouse influenza model. **A)** Study design: mice were randomized into five groups (n = 9-10/group): vehicle control, isotype control, APRIL-treated, APRIL/BAFF-treated, or unvaccinated. Treatments were administered twice weekly for the duration of the study. Influenza vaccination was performed on day 28 and 49. On day 70 animals were challenged with a mouse adapted Cal09 influenza strain and monitored daily for clinical signs and survival until day 91. **B)** Kaplan–Meier survival analysis for each group is plotted across the 21-day observation period post viral challenge. Statistical significance was assessed by log-rank test (Mantel-Cox) using GraphPad Prism. ns, not significant; *p < 0.05; **p < 0.01.

### Divergent effects of APRIL-only and dual APRIL/BAFF inhibition on serology, B-cell populations, and immunization

We next examined whether the survival differences observed in the challenge model correspond to measurable changes in humoral and cellular immunity. To this end, we profiled serum immunoglobulin isotypes (IgA, IgG, IgM) and performed cellular phenotyping of key immune compartments, including mature B cells, germinal center (GC) B cells, T follicular helper (Tfh) cells, and antibody-secreting cells (ASCs), in naive mice receiving repeated twice-weekly dosing with one of four treatments: APRIL inhibitor (4540), dual APRIL/BAFF inhibitors (POV or ATA, a recombinant protein with the same amino acid sequence as atacicept, produced internally; see Methods), or isotype control (MOTA). This approach allowed us to assess whether each mode of inhibition differentially affects antibody production, B-cell maturation, and downstream effector populations in the absence of antigenic challenge. Serological profiling revealed that both APRIL-only and dual APRIL/BAFF inhibition reduced serum IgA by ≥60% (Figure 2A), consistent with observations of IgAN patients treated with these inhibitors showing comparable IgA reduction^26, 33^. However, the two approaches diverged considerably in mice in their effects on other isotypes (Supplementary Figure S2). APRIL-only inhibition had a minimal impact on IgM (∼20% reduction) and a moderate effect on IgG (∼50% reduction), whereas dual APRIL/BAFF inhibition suppressed both IgM and IgG more broadly, with reductions of ≥70% and ≥60%, respectively. These patterns align with prior mouse studies in which APRIL-only blockade or genetic knockout sharply reduces IgA while sparing IgG and IgM to a greater degree than dual blockade^34, 35^. A similar qualitative divergence has also been observed in IgAN patients, where repeated administration of povetacicept produced sustained immunoglobulin reductions of approximately 40–50% in IgG and 80-90% in IgM through 48 weeks, compared with approximately 20-30% and 60-70%, respectively, reported with APRIL-selective inhibition^26, 36^. Collectively, these data indicate that selective APRIL blockade can substantially reduce IgA while largely preserving the IgG compartment, whereas dual inhibition, particularly with a potent agent such as POV, suppresses all major isotypes more broadly.

**Figure 2.**
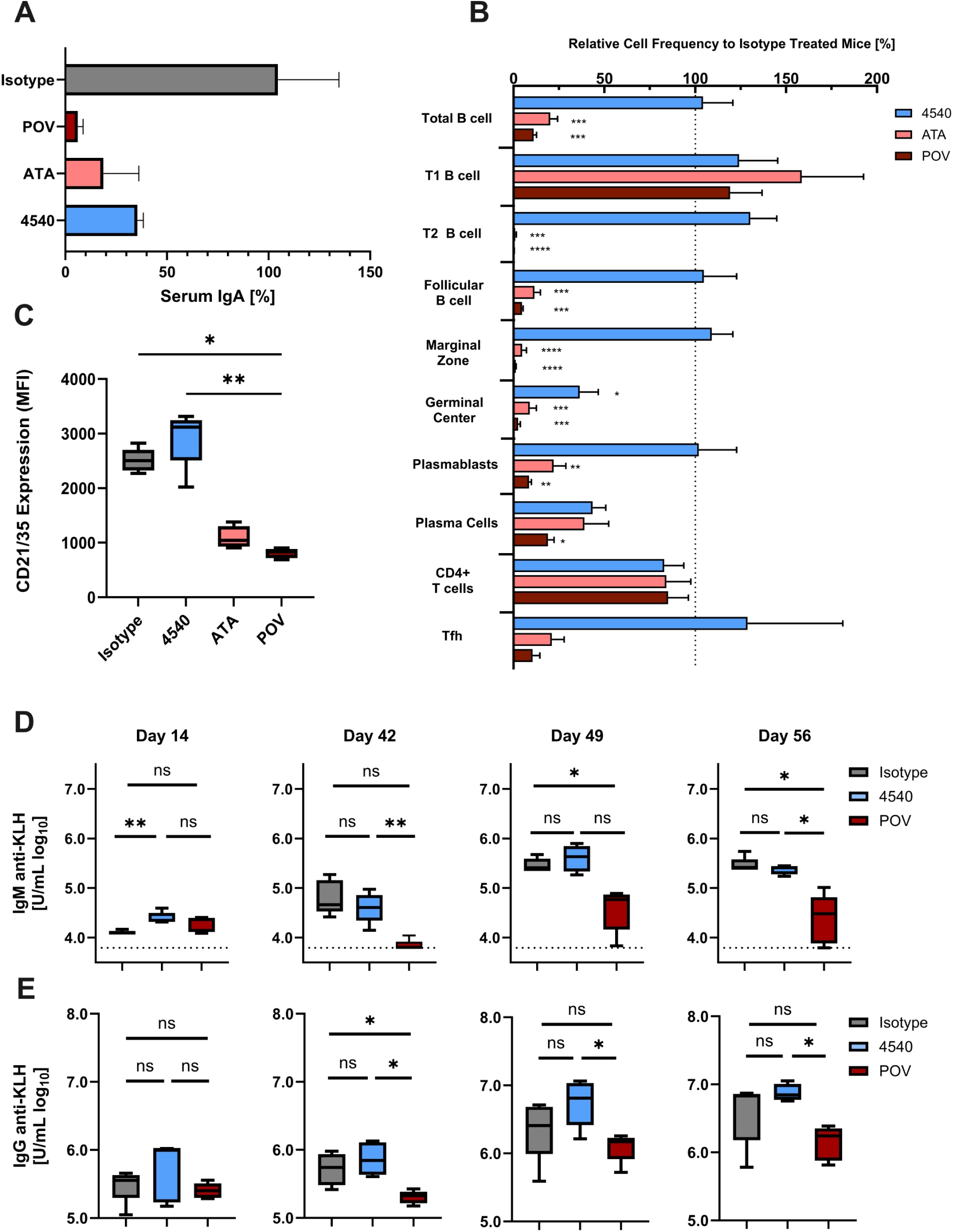
APRIL and APRIL/BAFF inhibition similarly reduce serum IgA but differentially impact splenic B-cell populations and immune response. A) Serum IgA levels measured at week 8 in mice treated with APRIL inhibitor (4540), APRIL/BAFF dual inhibitors (POV or ATA), or isotype control. IgA titers are shown relative to pre-treatment baseline and normalized to the mean isotype control measurement. Bars represent mean ± SEM. Statistical analysis was performed using Welch’s ANOVA with Dunnett’s T3 multiple-comparisons test. **B)** Flow cytometric analysis of splenic B- and T-cell subsets. Frequencies are expressed relative to the corresponding subset frequency in isotype-treated mice. Bars represent mean ± SEM. Statistical analysis was performed using one-way ANOVA with Tukey’s multiple-comparisons test. Significance is indicated as ns, not significant; *p < 0.05; **p < 0.01; ***p < 0.001; ****p < 0.0001 **C)** CD21/35 expression on splenic follicular B cells, shown as median fluorescence intensity (MFI). Boxes indicate the interquartile range, center lines indicate the median, and whiskers denote the range. Statistical analysis was performed using the Kruskal-Wallis test with Dunn’s multiple-comparisons test. Significance is indicated as ns, not significant; *p < 0.05; **p < 0.01; ***p < 0.001; ****p < 0.0001. KLH-specific serum IgM **D)** and IgG **E)** responses were measured over time in mice immunized with KLH and subsequently treated with isotype control (Mota), APRIL inhibitor (4540), or APRIL/BAFF dual inhibitor (POV). Mice were immunized on day 0 and day 42, and inhibitor treatment was initiated on day 14 at 10 mg/kg twice weekly and continued through day 56. Serum antibody titers were assessed on days 14, 42, 49, and 56. Day 14 represents the pre-treatment time point following KLH prime, day 42 represent responses after treatment during the primary immune response, and day 49 and 56 represents the recall response after KLH boost. Upper panels show KLH-specific IgM titers and lower panels show KLH-specific IgG titers. Box plots summarize group distributions, with individual animals overlaid. Dotted line represents the lower limit of quantification. Statistical comparisons between groups at each time point are indicated on the plots. One-way ANOVA statistical analysis was performed for IgM day 14, 42, 49 and IgG Day 42 and 49 using Brown-Forsythe and Welch with post-hoc Dunnett’s T3. IgM day 56 and IgG day 14 and 56 were analyzed using Kruskal-Wallis with post-hoc Dunn’s tests *p = <0.05, **p = <0.01.

We next performed cellular phenotyping of immune populations from the spleens of inhibitor-treated mice to determine whether the serological differences described above reflect corresponding changes in the underlying B cell and T cell compartments (Supplementary Figure S3). Repeated dosing with the anti-APRIL antibody 4540 did not significantly reduce total splenic B-cell numbers relative to isotype control, whereas both dual APRIL/BAFF inhibitors led to a greater than 80% depletion of total B cells (p < 0.001, Figure 2B). This divergence extended to specific B-cell subsets. Consistent with the known dependence of activated B cells, plasma cells, and IgA class switching on APRIL signaling^17, 37^, APRIL-only inhibition selectively reduced germinal center B cells and plasma cells but left other B-cell lineage subsets largely unaffected. Dual APRIL/BAFF inhibition, by contrast, caused significant reductions across nearly all B-cell subsets, including follicular B cells, germinal center B cells, and antibody-secreting cells, with the notable exception of T1 transitional B cells. The sparing of T1 cells is consistent with prior work demonstrating that B cells can mature in the bone marrow and migrate to the spleen as T1 cells independent of BAFF but require BAFF signaling for further developmental progression^38^, effectively creating a developmental bottleneck under dual blockade. The severity of B-cell depletion in dual-inhibitor-treated mice was also evident in spleen size: average total splenocyte counts were approximately 47-50 x10^6^ cells for APRIL-only and isotype control groups, respectively, compared to 21–28 x 10^6^ cells for the dual inhibitor groups. These results demonstrate that while both APRIL-only and dual APRIL/BAFF inhibition effectively reduce IgA, dual blockade induces broad depletion of most mature B-cell populations, whereas selective APRIL inhibition preserves the majority of the B-cell compartment.

The impact of dual inhibition was not limited to the B-cell lineage. While neither APRIL-only nor dual APRIL/BAFF inhibition significantly altered total T cell abundance relative to isotype control (Figure 2B), dual blockade, but not APRIL-only treatment, significantly reduced Tfh cells (Figure 2B), consistent with a reported role for BAFF in promoting Tfh development^39^. As Tfh cells are essential for supporting germinal center reactions and the generation of high-affinity antibodies, their loss represents an additional layer through which APRIL/BAFF inhibition may compromise adaptive immunity beyond direct B-cell depletion.

Beyond changes in cell abundance, dual blockade was also associated with shifts in surface phenotype among surviving B cells. Follicular B cells from dual APRIL/BAFF-treated mice exhibited reduced expression of CD21/35 (CR2), a component of the B cell coreceptor complex, whereas this reduction was not observed with APRIL-only inhibition (Figure 2C). Because CD21 expression increases with B cell maturation and is associated with antigen responsiveness, this observation is consistent with the pattern of impaired B cell maturation seen across multiple readouts under dual blockade.

In light of the extent of B-cell depletion and Tfh reduction under dual blockade, we next asked whether these changes translate to impairment of antigen-specific immune responses. To test this directly, we immunized mice with KLH on days 0 and 42 and beginning two weeks after the initial immunization (day 14) treated mice twice weekly with either an APRIL inhibitor (4540), a dual APRIL/BAFF inhibitor (POV), or isotype control (MOTA) for the duration of the study. This design allowed us to assess the impact of each treatment on both recall of established immunologic memory and on boosting of an ongoing response, which represent two clinically relevant scenarios for patients on chronic therapy. Anti-KLH IgM titers were significantly reduced in dual-inhibitor-treated mice both before (Day 42) and after (Days 49 and 56) the KLH boost, whereas APRIL-only treatment had no effect on anti-KLH IgM at any time point (Figure 2D). A similar pattern was observed for anti-KLH IgG: dual inhibition led to a trend toward reduced titers relative to isotype control and significantly lower titers at Days 42, 49, and 56, while APRIL-only treatment had no discernible effect on anti-KLH IgG levels (Figure 2E). These results demonstrate that dual APRIL/BAFF inhibition impairs both the maintenance and continued development of antigen-specific humoral responses, whereas APRIL-only inhibition preserves antigen-specific immunity even under sustained treatment.

Taken together, these results indicate that both modes of inhibition effectively reduce IgA, the pathologically relevant isotype in IgAN, but strongly diverge in their broader immunological impact. Selective APRIL inhibition largely preserves humoral and cellular immune function, whereas dual APRIL/BAFF blockade drives widespread disruption. Having demonstrated this pattern in mice, we next asked whether this divergence in B-cell impacts extends to human biology.

### Dual but not APRIL-selective inhibition reduces human B-cell survival in vitro

To determine whether the differential effects in mice extend to human B cells, we evaluated the contributions of APRIL and BAFF to primary human B-cell survival in vitro using an established assay^40, 41^. Human B cells purified from peripheral blood were first cultured for 3 days in the presence of IL-2 and CD40L, then washed and maintained for five additional days in the presence of APRIL, BAFF or both cytokines before viability was measured (Figure 3A). BAFF promoted B-cell survival relative to cytokine-deprived conditions, whereas APRIL had no appreciable effect. Co-stimulation with both cytokines did not enhance survival beyond BAFF alone, indicating BAFF is the dominant survival factor in this system.

**Figure 3.**
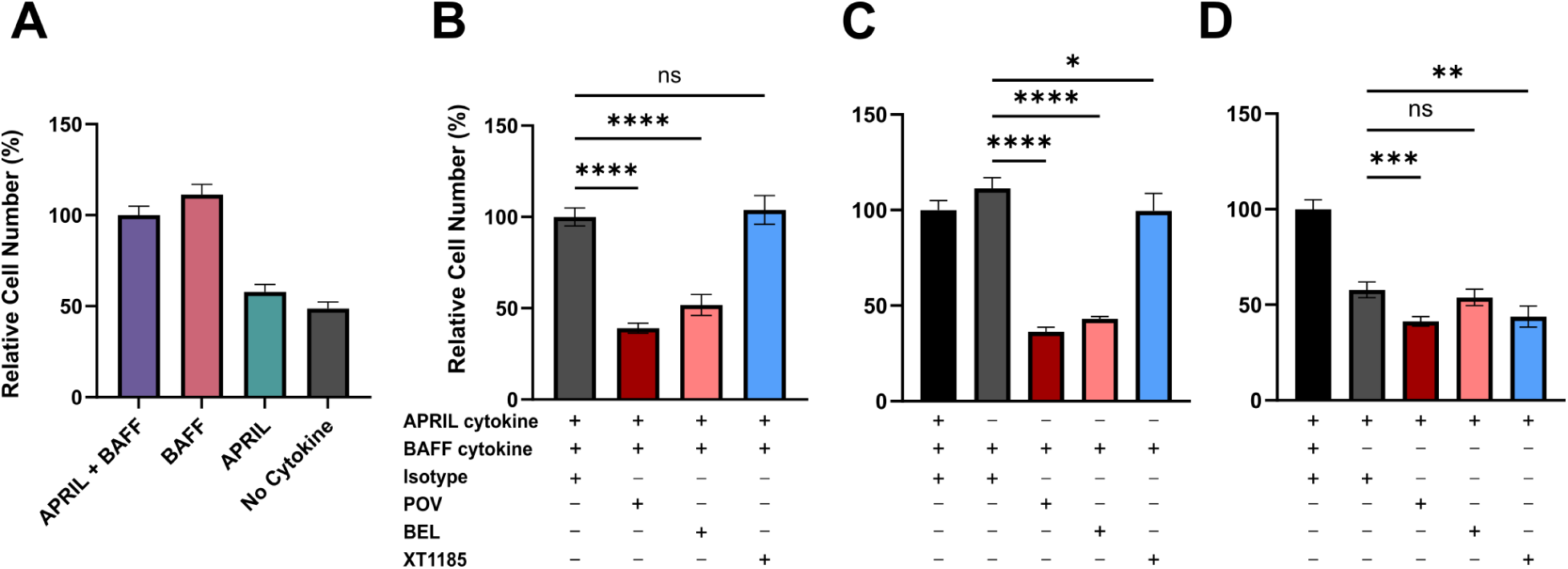
APRIL, BAFF, and dual APRIL/BAFF cytokine inhibitors differentially regulate survival of human B cells in culture. **A)** Human B-cell viability was measured 5 days after stimulation with BAFF, APRIL, or APRIL + BAFF. Viability was quantified using the CellTiter-Glo luminescent assay as a measure of metabolically active cells and normalized to the APRIL + BAFF stimulation condition. **B–D)** Human B cells were cultured under stimulation with APRIL + BAFF (B), BAFF alone (C), or APRIL alone (D) in the presence of APRIL- and BAFF-targeting inhibitors POV (dual APRIL/BAFF inhibitor), BEL (BAFF-selective inhibitor), or XT1185 (APRIL-selective inhibitor). Cell viability was measured after 5 days by CellTiter-Glo and normalized to the isotype control for the APRIL + BAFF stimulation condition. Data are shown as mean ± SD. Statistical significance was determined by one-way ANOVA with Dunnett’s multiple-comparison test relative to the corresponding isotype control. ns, not significant; *p < 0.05; **p < 0.01; ***p < 0.001; ****p < 0.0001.

We next assessed how selective or dual cytokine blockade influences B cell survival. Across stimulation conditions containing BAFF, either alone or in combination with APRIL, treatment with a dual APRIL/BAFF inhibitor or a BAFF-only inhibitor markedly reduced cell survival, whereas selective APRIL inhibition had limited impact (Figure 3B,C). Under APRIL-only culture conditions (Figure 3D), selective APRIL blockade modestly reduced viability to a level comparable to dual inhibition, while BAFF inhibition had minimal effect, consistent with APRIL providing a limited survival contribution to the majority of the B cell population.

Collectively, these findings are consistent with the in vivo observations in mice and support a model in which APRIL-targeted therapies can reduce IgA without substantially compromising B cell survival, a distinction with relevance for chronic dosing in diseases such as IgA nephropathy.

### Single-cell multi-omics characterization of differential impacts on splenic immune populations

To more deeply characterize the cellular and molecular consequences of each mode of inhibition, we applied two complementary multi-omic approaches to the spleen. First, we employed single-cell CITE-seq, which yields paired transcriptomic and surface proteomic data from 129 surface proteins per cell, enabling a direct comparison with our flow cytometric findings while also revealing gene expression changes within surviving populations. Second, recognizing that the organized architecture of the spleen, and particularly the structure of B-cell follicles and germinal centers, is important for orchestrating adaptive immune responses, we applied single-cell-resolution in situ spatial transcriptomics (10x Genomics Xenium), with a 5,000-gene panel, to resolve immune cell subsets within their native tissue niches.

To characterize the impact of each mode of inhibition on splenic immune populations at single-cell resolution, we performed CITE-seq on splenocytes from mice chronically dosed with APRIL inhibitor, dual APRIL/BAFF inhibitor, or isotype control (Methods). Cell type annotation using marker genes and surface proteins identified 29 distinct cell types (Figure 4A; Supplementary Figure S4), including several B-cell subpopulations relevant to APRIL and BAFF biology. Unsupervised clustering of the B cell and ASC compartment identified 8 distinct populations, which were annotated using canonical surface protein markers from the CITE-seq antibody panel (Figure 4C; Supplementary Figure S5): transitional B cells (T1, T2) were identified by CD93 expression with progressive IgD, CD23 (FCER2A) and CD21 (CR2) acquisition; follicular B cells by high IgD, CD21, and moderate CD23 (FCER2A); marginal zone B cells by high CD21, IgM, CD1d1, and low CD23; germinal center B cells by Ki67 and AICDA expression; B-1 cells by CD43 (SPN) expression with high IgM with low B220 (PTPRC), IgD and CD93; and ASCs by high JCHAIN and SDC1 (CD138) with low CD19. We also identified a distinct cluster that expressed canonical markers of follicular B cells but exhibited a unique gene expression signature, including elevated gene expression of *Runx1*, a transcription factor reported to influence BCR responsiveness^42^. We annotated these cells as Runx1^hi^ follicular B cells (Figure 4B,C).

**Figure 4.**
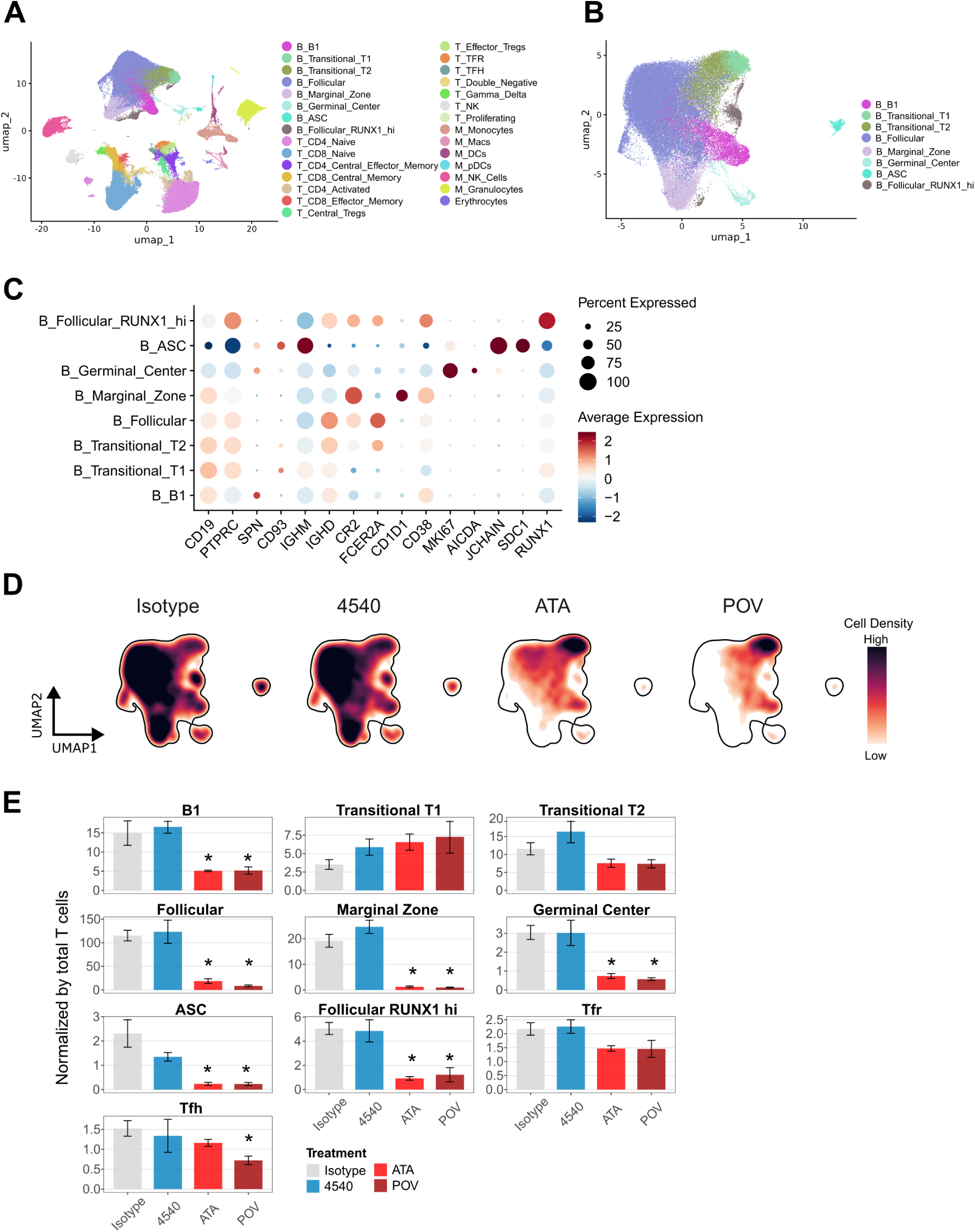
CITE-seq confirms treatment-dependent changes in splenic B cell populations. **A)** Annotated UMAP from RNA expression profiles of spleen cells (141,811 total cells) from treated and control mice (n=3-5 per group). **B)** B cells, including ASCs, represented eight distinct transcriptional subsets and annotated based on marker genes and proteins. **C)** Selected proteins were visualized across B cell clusters. The dot color indicates the normalized average protein expression and size reflects the percentage of cells within each cluster expressing the marker. **D)** UMAP coordinates were converted to kernel density estimates for each treatment group to support regional changes in B cell composition. The outlined area indicates expected cell locations. **E)** Quantification of B cell subsets’ abundance normalized to total T cells per sample. Significance was determined using an unsigned Wilcoxon unsigned rank-sum test in which all three treatments were compared against isotype (*p<0.05).

We first assessed how B-cell abundance was impacted by each treatment: antibody 4540, ATA, POV, or an isotype control (Figure 4D-E). Consistent with our flow cytometric phenotyping, CITE-seq data showed that dual APRIL/BAFF inhibition led to substantial reductions in all B cell and ASC subsets except T1 transitional B cells. APRIL-only inhibition, by contrast, led only to a partial reduction in ASCs, leaving other B-cell subsets largely unaffected. Given our earlier observation of Tfh cell impacts by flow cytometry, we also inspected this population in the CITE-seq data and confirmed that APRIL-only inhibition had no appreciable effect on Tfh abundance, while dual inhibition reduced Tfh cells, with the more potent inhibitor POV producing a greater reduction. Dual APRIL/BAFF inhibitors, but not APRIL-only, showed a trend towards a reduction in regulatory T follicular (Tfr) cells. Collectively, these single cell multi-omic results are consistent with the flow cytometry results, confirming that dual APRIL/BAFF blockade broadly depletes B-cell populations and Tfh cells, while APRIL-only inhibition has a far more circumscribed impact.

### Spatial transcriptomics reveals collapse of follicular architecture under dual APRIL/BAFF inhibition

Having characterized splenic cellular composition by CITE-seq, we next applied in situ spatial transcriptomics to resolve how these abundance changes manifest within tissue architecture. Because this platform operates on fixed tissue sections, it also enabled recovery of stromal and endothelial populations that are underrepresented in dissociated single-cell approaches but play important roles in organizing lymphoid tissue structure. Spatial data from spleen were obtained for PBS control (n=1), isotype control (n=2), 4540- (n=3), ATA- (n=2), and POV-treated (n=3) mice.

Unsupervised clustering identified 15 populations spanning immune and non-immune lineages (Supplementary Figure S6). Within the B-cell lineage, we identified three major populations: CR2^hi^ B cells (enriched in follicular B cells), CR2^lo^ B cells (enriched in transitional, marginal zone, and B-1 cells), and germinal center B cells, alongside ASCs. Clusters with mixed lineage markers consistent with known segmentation artifacts of ISH-based platforms^43^ were identified and excluded from downstream analyses (Supplementary Methods).

In control spleens, the localization of annotated cell types revealed the expected follicular architecture: regions enriched in CR2^hi^ B cells surrounding central T cell zones, smaller regions of concentrated GC B cells indicating the presence of germinal centers, CR2^lo^ B cells enriched at the follicle periphery and in non-follicular regions, ASCs clustered outside the follicles, and supportive stromal and endothelial cells in non-follicular regions (Figure 5A and Supplementary Figure S7). Compared to this baseline, the effects of each inhibitor were readily apparent (Figure 5B-D). APRIL-only inhibition with 4540 did not significantly alter cell populations or the overall organization of follicular structures, aside from a potential reduction in ASCs. Dual APRIL/BAFF inhibition, by contrast, led to substantial decreases in CR2^hi^ B cells, GC B cells, and ASCs, and these losses were not evenly distributed or subtle in their structural consequences. Follicle-like structures showed a striking depletion of B cells and became composed primarily of T cells, visually underscoring the extent of immune disruption.

**Figure 5.**
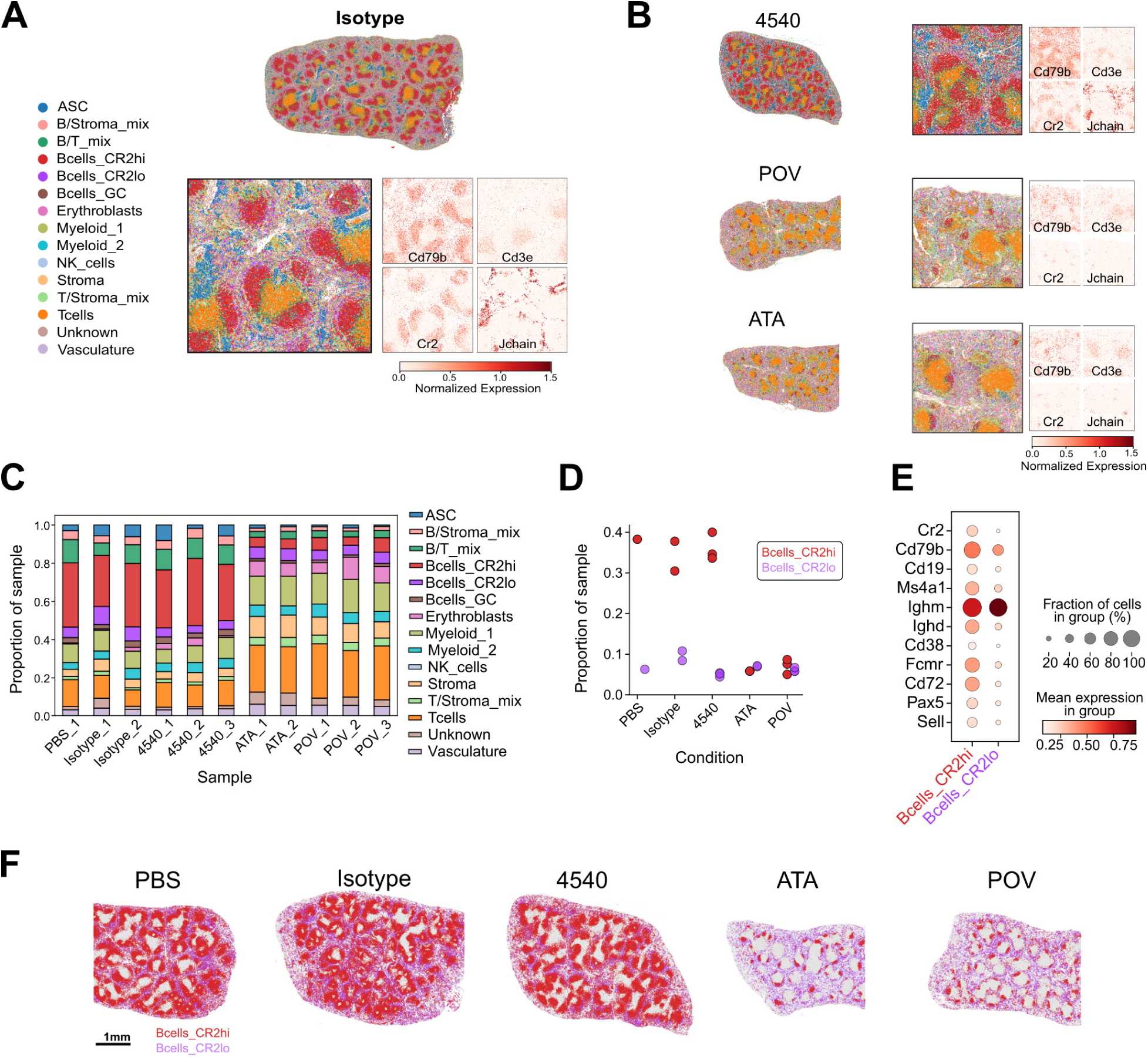
Spatial transcriptomics illustrates CR2^hi^-specific B cell depletion in mice treated with APRIL/BAFF dual inhibitors, compared to APRIL inhibition alone. A, B) Representative spleen samples of mice treated with isotype, anti-APRIL (4540) or anti-APRIL/BAFF (ATA, POV) subject to Xenium spatial transcriptomics analysis. Cell types were annotated using validated lineage markers, highlighting follicular B and T cell zones. Each marker represents log normalized expression. **C)** Cell type proportions quantified as the fraction of each cell type relative to total cells detected per sample. **D)** B cell proportions of CR2^h^^i^ and CR2^lo^ expressing subsets compared across treatment groups. **E)** Representative genes differentiating CR2^hi^/CR2^lo^ B cell subsets in tissue. Color intensity represents the mean log-normalized expression per cell types, and the size represents the percent of cells with positive expression. **F)** Visualization of CR2^hi^ and CR2^lo^ B cells in representative mouse spleens across experimental conditions. Remaining cell types are represented in light gray.

Characterization of B-cell marker gene expression confirmed the identity of our spatial clusters: CR2^lo^ B cells were enriched for high *Igm* expression, supporting their annotation as transitional and marginal zone B cells, while CR2^hi^ B cells showed greater *Igd* expression consistent with follicular B-cell enrichment (Figure 5E). Mapping these populations across the entire spleen demonstrated a consistent loss of CR2^hi^ (follicular-enriched) B cells across all regions of the organ under dual blockade (Figure 5F), indicating that the depletion is not confined to particular splenic zones but reflects a global collapse of the follicular B-cell compartment. Taken together, these spatial data show that dual APRIL/BAFF inhibition goes beyond reducing B-cell numbers; it disrupts the follicular structures which are organized to support adaptive humoral responses. These architectural changes prompted us to investigate the underlying molecular mechanisms, including the chemokine networks and gene programs that govern germinal center formation and function.

### Dual but not APRIL-only inhibition disrupts germinal center architecture and molecular programs supporting GC function

The architecture and cell composition of lymphoid follicles underpins the interaction of B cells with T cells to form germinal centers, which are the structures responsible for generating high-affinity, class-switched antibodies and durable B-cell memory. Given the abundance changes in CR2^hi^ B cells and GC B cells, we investigated whether the molecular infrastructure supporting GC formation and lymphocyte homing was also affected. Using key GC B-cell markers including *Aicda*, *Bcl6*, *Cxcr4*, and *Mki67*, we identified spatial regions with concentrated expression of these markers, indicating the presence of germinal centers within B cell-rich follicular zones (Figure 6A). Quantification of total GC number per spleen (Methods) revealed a near-total loss of germinal centers in spleens of dual APRIL/BAFF inhibitor-treated mice, while APRIL-only inhibition had minimal impact on GC numbers (Figure 6B).

**Figure 6.**
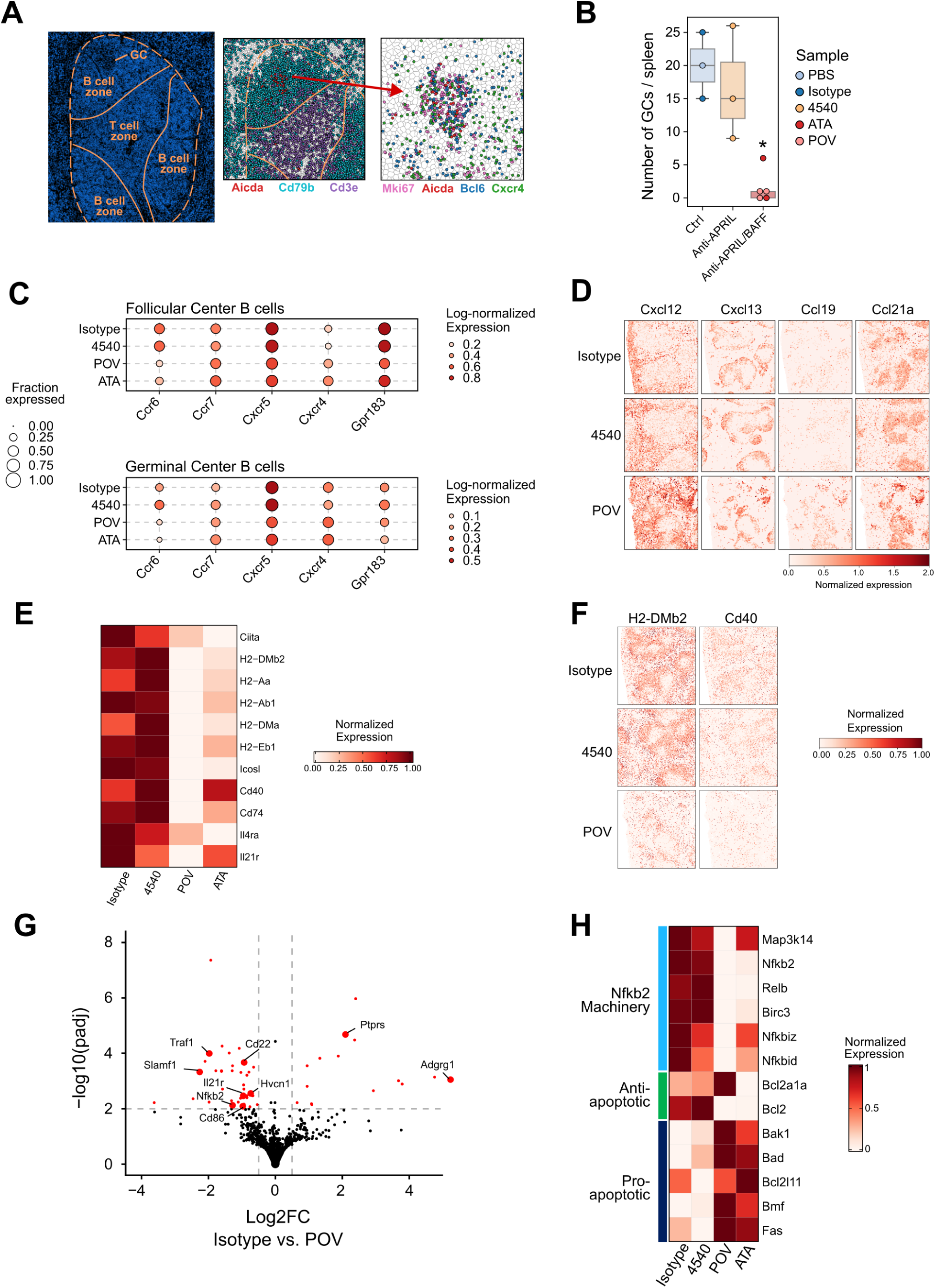
Dual (APRIL/BAFF) inhibition disrupts germinal center (GC) formation, antigen presentation, and B cell survival in the spleen. A) Representative germinal center reaction shown by DAPI was supported by co-expression of GC-enriched markers: *Mki67*, *Bcl6*, and *Aicda*, with each point representing an individual transcript. B) Germinal centers were quantified per spleen and compared by treatment groups using two-sided exact permutation tests compared to sample controls. The anti-APRIL group was not significantly different from control (p = 0.80), whereas the anti-APRIL/BAFF was (p = 0.036). **C)** Quantification of average log-normalized expression values for chemokine receptors with known roles in GC formation across treatment groups. Data are derived from single-cell expression results. **D)** Non-B cell associated chemokines represented in representative spleen regions across treatment groups. Values are log-normalized expression. **E)** Heatmap of key genes mediating B cell–Tfh cell interactions in GC B cells. Values are derived from mean single-cell expression data and scaled from 0 to 1. **F)** Spatial expression of MHC class II-associated genes across conditions in representative spleen regions. Values are log-normalized. **G)** Differential expression analysis performed on pseudobulked GC B cells derived from single-cell expression data. Significance was determined using the DESeq2 likelihood ratio test. **H)** Heatmap of B cell survival pathway genes across treatment groups.

The organization of splenic microarchitecture, including the formation and maintenance of GCs, depends on a coordinated network of chemokines and their receptors expressed by lymphocytes, stromal cells, and endothelial cells. We therefore investigated whether dual inhibition disrupts these chemokine networks. In our CITE-seq data, which provides richer gene expression information for lymphocyte populations, we examined expression of key homing receptors on follicular and GC B cells (Figure 6C) as well as Tfh and Tfr cells (Supplemental Figure S8). *Ccr6* and *Cxcr5*, both of which play important roles in B-cell entry into and positioning within the GC, showed reduced gene expression in follicular and GC B cells under dual inhibition but not under APRIL-only conditions (Figure 6C). CCR6 has been shown to mark GC memory B-cell precursors in the light zone and to modulate the kinetics of GC entry^44, 45^, while CXCR5 is the cognate receptor for CXCL13 and is essential for B cell and Tfh cell migration into follicles, with *Cxcr5*-deficient mice showing defective formation of primary follicles and GCs^46^. The downregulation of both receptors under dual blockade thus suggests impaired capacity for follicular homing and GC participation among the remaining B cells. Conversely, *Cxcr4*, which mediates centroblast localization to the GC dark zone where proliferation and somatic hypermutation occur^46^, showed increased gene expression under dual inhibitor but not APRIL-only conditions (Figure 6C), potentially reflecting a compensatory shift or altered GC B-cell dynamics in the context of substantial B-cell depletion.

A notable limitation of our CITE-seq dataset is the underrepresentation of stromal and endothelial cells, likely due to poor survival of these populations through tissue processing and cryopreservation. However, these non-hematopoietic populations are major producers of chemokines that organize lymphoid tissue architecture. We therefore leveraged our spatial transcriptomics data, which, being derived from fixed tissue, retains these populations, to characterize chemokine expression from stromal and endothelial sources.

*Cxcl12*, *Cxcl13*, *Ccl19*, and *Ccl21a* all showed increased expression in spleens from dual inhibitor-treated mice relative to isotype control and APRIL-only treatment (Figure 6D). These chemokines collectively orchestrate lymphoid tissue organization: CXCL13, produced by follicular dendritic cells and stromal cells, is the primary chemoattractant directing CXCR5+ B cells and Tfh cells into follicles^47^ and has been reported to synergize with BAFF in supporting follicle organization^48^. CXCL12, produced by reticular cells in the GC dark zone, works in concert with CXCL13 to establish the dark zone–light zone polarity essential for affinity maturation^46, 49^. CCL19 and CCL21, the ligands for CCR7, direct T cells and dendritic cells to appropriate positions within the white pulp, with their absence leading to disrupted T and B-cell compartmentalization^50^. Notably, the upregulated chemokines also showed a more concentrated and spatially localized expression pattern under dual inhibitor conditions. In the setting of profound B-cell depletion, the elevated expression of follicular and T-zone chemokines likely reflects compensatory stromal remodeling, in which the tissue microenvironment attempts to recruit lymphocytes to niches that have been vacated. Such active stromal responses have been observed in other contexts of lymphocyte depletion^51–53^ and suggest that dual APRIL/BAFF inhibition induces not only cell loss but disrupts the stromal–lymphocyte crosstalk that maintains splenic architecture.

We next investigated whether antigen presentation and costimulatory programs were differentially affected. While APRIL-only inhibition led to minimal changes in the expression of antigen presentation genes (H2 family, *Ciita*), dual APRIL/BAFF inhibition showed substantial expression reduction of these genes in GC B cells (Figure 6E) and in follicular B cells (Supplementary Figure S8). Key B-cell costimulatory molecules including ICOSL, CD40, and CD74, as well as supportive cytokine receptors IL4RA and IL21R that facilitate productive GC B cell–Tfh interactions, were also broadly downregulated under dual inhibition with POV and ATA, whereas 4540 dosing had minimal effect on these gene programs (Figure 6E). In particular, CD40 expression on B cells, but not on dendritic cells, has been shown to be required for generating antigen-specific GC B cells and Tfh cells as well as high-affinity class-switched antibody production^54^. Similar patterns were observed in the spatial transcriptomics data (Figure 6F and Supplementary Figure S9). Together, these findings suggest that dual APRIL/BAFF blockade not only depletes the cells involved in GC formation and activity but also impairs the molecular machinery required for productive B cell–T cell interactions in the remaining population, which is consistent with the impaired antigen-specific responses observed in the KLH immunization model.

Finally, to further elucidate the mechanistic basis for these differential effects, we performed pseudobulk differential expression analysis in GC B cells comparing APRIL-treated, dual APRIL/BAFF-treated and isotype control mice. This analysis revealed minimal differentially expressed genes for APRIL-only inhibition with 4540 (Supplementary Figure S10), whereas dual inhibition with POV produced widespread transcriptional changes (Figure 6G). Among the most prominent was a coordinated reduction in components of the noncanonical NF-κB2 signaling pathway, including *Map3k14* (encoding NIK), *Nfkb2*, *Relb*, *Birc3*, *Nfkbiz*, and *Nfkbid* (Figure 6H). This pathway is the primary conduit through which BAFF-R delivers survival signals to B cells, with BAFF binding resulting in stabilizing NIK and enabling processing of p100 to p52 and nuclear translocation of RelB:p52 dimers^55, 56^. The reduced expression of this pathway is consistent with removal of BAFF-mediated tonic survival signaling under dual blockade. Downstream of this, GC B cells from dual-inhibitor-treated mice showed a shift in the apoptotic balance, with pro-apoptotic genes including *Bak1*, *Bad*, *Bcl2l11*, *Bmf*, and *Fas* upregulated, and a trend toward reduced expression of anti-apoptotic factors. Beyond survival programs, dual inhibition also significantly reduced expression of genes with established roles in GC function, including *Cd86* (costimulation during antigen presentation), *Il21r* (a cytokine receptor supporting GC formation and B-cell proliferation^57^), *Slamf1* (whose deletion leads to attenuated B-cell responses^58^), *Cd22* (important for memory B-cell formation^59^), and *Hvcn1* (a BCR-associated channel whose loss impairs class switching^60^).

Taken together, our single-cell and spatial transcriptomics analyses demonstrate that dual APRIL/BAFF inhibition disrupts splenic immune function at multiple levels, from GC structure and follicular chemokine organization to antigen presentation, costimulatory programs, and B-cell survival signaling. In contrast, APRIL-only inhibition leaves these programs largely intact. These mechanistic findings provide a molecular explanation for the divergent functional outcomes observed in both the influenza challenge and KLH immunization models and support that selective APRIL inhibition achieves IgA reduction without the broad immune compromise associated with dual APRIL/BAFF blockade.

## DISCUSSION

B-cell dysfunction is a hallmark of immune-mediated kidney diseases, including lupus nephritis, IgA nephropathy (IgAN), ANCA-associated vasculitis, and primary membranous nephropathy^8, 61^. B-cell development occurs through a highly regulated process that is dependent on several receptors and cytokines, many of which have been targeted by different therapeutic approaches. Depending on the disease, and the etiology of B-cell dysregulation within the disease, different approaches may prove more or less efficacious. For example, rituximab, an anti-CD20 B-cell depletion antibody, has demonstrated clinical efficacy in primary membranous nephropathy^62, 63^ but is less effective in IgAN^64^. These observations underscore the importance of matching therapeutic mechanism to disease biology.

IgAN is one such disease where systematic biochemical and mechanistic studies have identified a well-characterized pathogenic framework, which is captured in the four-hit model^1, 65^. Recent clinical data has suggested that targeting APRIL alone or in combination with BAFF may provide therapeutic benefit in IgAN^26, 66, 67^, potentially through modulation of one or more steps within the four-hit model. There is strong genetic and mechanistic support that APRIL is a key driver of the pathogenesis of IgAN^22, 23^, likely through mediating class switching within mature B cells and promoting plasma cell survival, thereby promoting production of Gd-IgA1 and potentially also supporting the autoreactive humoral responses directed against Gd-IgA1 autoantigen. By contrast, the role of BAFF in IgAN is less clearly defined, with some in vivo data supporting a role for BAFF in the disease^68^, whereas other data indicating it may not play a central role to the pathogenic process^69^. Of note, clinical data with BAFF inhibition alone in IgA nephropathy indicates limited protective effect^25^.

Although direct head-to-head clinical comparisons are not available, late-stage clinical data suggest that both APRIL-selective and dual APRIL/BAFF inhibition can modulate key disease-associated biomarkers in IgAN, including IgA-related measures, proteinuria, and eGFR^26, 33, 66, 67^. However, there is not yet clear evidence that the addition of BAFF blockade yields a qualitatively greater benefit on these biomarkers or clinical outcomes. In the absence of such a benefit, the breadth of downstream immune perturbation becomes a central consideration, particularly because treatment of IgAN is likely to be chronic. Our phenotypic and mechanistic data, together with prior findings^39^, support the conclusion that dual APRIL/BAFF inhibition produces broader effects than APRIL-only inhibition on B-cell survival, chemokine programs, antigen-presentation capacity, and germinal center structure and function (Figure 7). Conceptually, these findings support a distinction between selective modulation of the pathogenic IgA axis and broader suppression of humoral immunity.

**Figure 7.**
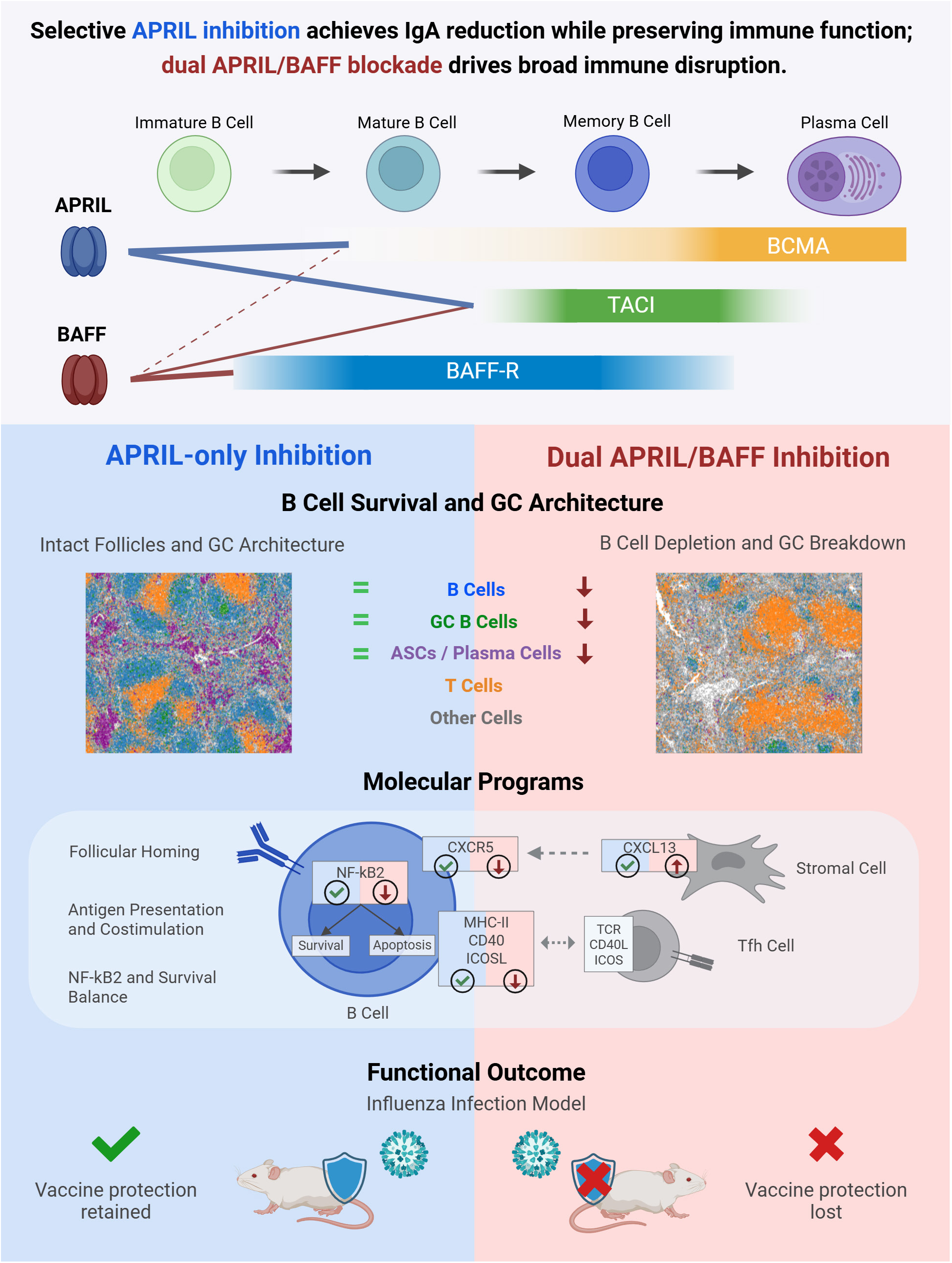
Summary of differential impact of APRIL/BAFF on B cell homeostasis and GC disruption.

In our studies, one consequence of dual APRIL/BAFF inhibition is a reduced serological response to vaccination which, in the case of an influenza challenge model, resulted in substantial loss of a protective immunity. This functional effect was concordant with the underlying cellular phenotype from our inhibitor-only studies, in which dual APRIL/BAFF inhibition was associated with lower immunoglobulin levels, reduced total B cells, fewer germinal center B cells, and fewer plasma cells. In contrast, APRIL-only inhibition had substantially less impact on these broader immune compartments. These findings suggest that the addition of BAFF blockade extends the effects of treatment beyond pathways most directly implicated in pathogenic Gd-IgA production and into mechanisms required for maintenance of humoral immune competence. Analysis of single cell and spatial transcriptomic data sets provides further mechanistic insight into pharmacologic effects of APRIL and APRIL/BAFF inhibition. These analyses indicated that dual inhibition altered pathways linked to B-cell survival, maturation, and antigen responsiveness. For example, NF-kB and LTB-associated programs are downregulated in T1 cells consistent with impaired progression through normal B-cell maturation states. In addition, expression of genes associated with B-cell survival and function (e.g., *Nfkb2*, *Icosl*, *Bcl2*, and *Ltb*) was reduced under dual inhibition, whereas pro-apoptotic programs were increased in T2 cells. Key transcription regulators including *Irf4*, *Bcl6*, and *Ciita*are impacted under dual inhibition, which govern responses to antigen challenge. Because IRF4 and BCL6 are central to germinal center dynamics and plasma cell differentiation^70, 71^, and CIITA is a master regulator of MHC class II expression^72^, these changes are consistent with reduced antigen-driven B-cell function and altered maturation. Taken together, the transcriptomic data support the broader phenotypic observation that dual APRIL/BAFF inhibition shifts the B-cell compartment toward less mature and less functionally responsive states.

Several lines of evidence support the translational relevance of these findings to humans. Human genetics provide the most direct link: homozygous deletion affecting *TNFRSF13C* (encoding BAFF-R) arrests B-cell development beyond the immature/transitional stage and is associated with agammaglobulinemia and impaired humoral responses^73, 74^, while repeated administration of atacicept in non-human primates produces marked reductions in immunoglobulins, particularly IgM, together with loss of total and mature B-cell populations^75^. By contrast, APRIL deficiency due to mutation in *TNFSF13* has been associated with reduced immunoglobulin levels and no clear deficiency in B-cell populations^76^, a phenotype more aligned with the narrower effects of APRIL blockade observed here. This genetic prediction is confirmed by clinical pharmacodynamic data: repeated administration of sibeprenlimab to patients with IgA nephropathy over 12 months preserved peripheral blood IgA+, IgM+, and inferred IgG+ memory B-cell populations with no clinically meaningful depletion compared to placebo (Supplementary Figure S11). These findings are consistent with the preservation of total B-cell and memory B-cell populations reported in the ENVISION COVID-19 substudy (McCafferty et al., manuscript under review). Although congenital deficiency and pharmacologic inhibition are not equivalent, the convergence across mouse, non-human primate, human genetic, and clinical datasets supports the conclusion that BAFF pathway inhibition has a broader impact on global B-cell homeostasis than APRIL-selective blockade.

Clinical observations in other autoimmune settings provide additional, although indirect, support for our interpretation. BAFF-pathway inhibition in patients has been associated with reduced vaccine responsiveness, including diminished responses to SARS-CoV-2 mRNA vaccination, consistent with impaired B-cell priming and effects on circulating T follicular helper cells^30^. Prolonged belimumab treatment in SLE has also been associated with marked reductions in circulating B-cell subsets, with naïve, memory, activated, and plasmablast/plasma cell populations decreased by approximately 40–99%^77^. Similarly, dual APRIL/BAFF inhibition with telitacicept has been reported to reduce total B cells, transitional B cells, naïve B cells, and short-lived plasma cells in lupus patients^78^. Reports of more severe COVID outcomes in lupus patients receiving telitacicept^79^, and hypogammaglobulinemia observed in 5-12% of IgAN patients with increasing doses of povetacicept in early-phase studies^33^, are directionally consistent with the broader immunologic footprint seen in our preclinical studies.

This study has several limitations. All in vivo experiments were conducted in wild-type mice rather than in a disease-relevant model of IgAN and therefore do not directly address how these differential immunological effects manifest in the context of Gd-IgA1 production, mesangial deposition, or proteinuria. In addition, the multi-omic and spatial transcriptomic analyses were focused on the spleen and do not capture the mucosal immune compartment. Given the role of gut-associated lymphoid tissue in IgA class switching and in the initiation of pathogenic IgA responses in IgAN, the effects of selective versus dual inhibition on mucosal B-cell populations and local IgA production represent an important area for future investigation.

In summary, our results support a model in which APRIL-selective inhibition more narrowly targets biology relevant to the pathogenic IgA axis in IgAN, whereas concomitant BAFF inhibition extends the pharmacologic effect to broader aspects of B-cell maturation, germinal center organization, and humoral immune competence. This distinction may be especially important in a chronic disease setting, where treatment duration is likely to be prolonged and preservation of baseline immune function is a meaningful component of therapeutic value. If disease-relevant IgA biomarkers can be equivalently reduced without BAFF co-inhibition, then minimization of broader immune perturbation becomes a key clinical consideration. Our findings collectively support the view that APRIL-only inhibition may offer a more favorable therapeutic index for long-term treatment of IgAN.

## METHODS

### Proteins

Anti-APRIL monoclonal antibody (mAb) 4540 was produced in-house using the Fv sequence disclosed in patent application US 2017/0145086 A1 and formatted with a human IgG1 Fc. Anti-APRIL mAb XT1185 was produced in-house using the variable-domain sequence of sibeprenlimab (Inxight Drugs, an NIH-curated database of approved, marketed, and investigational drugs; GHX28QZ7DD) and formatted with a human IgG1 Fc with immunosilencing mutations. Anti-APRIL/BAFF molecules POV and ATA were generated to match the sequences of povetacicept (Inxight Drugs, EN2U82XV65) and atacicept (Inxight Drugs, K3D9A0ICQ3), respectively. ATA was also purchased from MedChemExpress (HY-P99446). The anti-BAFF belimumab biosimilar was purchased from InvivoGen (hbaff-mab1). An isotype control mAb based on the sequence of motavizumab (Inxight Drugs, 50Y163LK8Q) was produced in-house or by GenScript or Icosagen. Proteins were expressed in Expi293 or CHO cells, purified by Protein A affinity chromatography, and further purified by preparative size-exclusion chromatography on a Superdex 200 Increase 10/300 column. Fractions corresponding to the main monomeric peak were pooled and dialyzed into 1× PBS.

### Naive Mouse Study: APRIL and APRIL/BAFF Inhibitor Dosing

Male C57BL/6J mice (6 weeks old; 19-25 g) were purchased from The Jackson Laboratory (Bar Harbor, ME, USA). In all studies, mice were randomly assigned to groups and received 10 mg/kg of the indicated inhibitor or control by intraperitoneal injection twice weekly. Blood was collected weekly by submandibular puncture, and terminal blood and spleens were collected at necropsy. Plasma was isolated and stored at -80°C until analysis. All animal procedures were conducted in accordance with the National Institutes of Health Guide for the Care and Use of Laboratory Animals and were approved by the Institutional Animal Care and Use Committee of Mispro Biotech Services.

For the 8-week study, mice were randomized to receive isotype control, 4540, ATA, or POV (n=5 per group at study start). One ATA-treated animal did not survive to the scheduled endpoint, resulting in n=4 for terminal analyses in that group. Spleens were processed into single-cell suspensions and cryopreserved in CryoStor CS10 medium (STEMCELL Technologies) prior to analysis. Flow cytometry was performed using splenocytes from n=5 isotype control, n=5 4540, n=5 POV, and n=4 ATA animals. The same splenocyte preparations were used for single cell CITE-seq, however, because of limited splenocyte availability in APRIL/BAFF inhibitor-treated groups, single-cell profiling was performed on n=5 isotype control, n=5 4540, n=4 POV, and n=3 ATA samples.

For the 4-week spatial transcriptomics study, mice were randomized to receive isotype control (n=2), 4540 (n=3), PBS vehicle (n=2), POV (n=3), or ATA (n=2). At 4 weeks after treatment initiation, spleens were collected, fixed intact in 10% neutral buffered formalin at 4°C for 144 hours, dehydrated through graded ethanol, cleared in xylene, and embedded in paraffin.

### Serology

Total serum IgA, IgM, and IgG measurements were performed using enzyme-linked immunosorbent assay (ELISA) kits (E99-103, E99-101, E99-131; Bethyl Laboratories) following the manufacturer’s instructions. Sera samples were minimally diluted 1:400 for IgA analysis, 1:1,000 for IgM, and 1:5,000 for IgG. Sera samples with high immunoglobulin concentrations were diluted further. For each timepoint, immunoglobulin values were interpolated from a standard curve generated with 4-parameter logistic curve fit and 1/Concentration2 weighting using Softmax Pro. Data was plotted and one-way ANOVA analyses were performed for the terminal sera timepoint comparing isotype control to each type of inhibitor using GraphPad Prism v10.4.1.

### Flow cytometry of splenic immune cells

Cryopreserved splenocytes from the 8-week dosing study were thawed and immunophenotyped by flow cytometry to quantify splenic B cell, T cell, and antibody-secreting cell populations. Samples were acquired on a Sony ID7000 flow cytometer using antibody panels prepared at empirically optimized concentrations for each clone and fluorochrome combination (Supplementary Table S1). Fluorescence minus one controls were used to guide gate placement.

Briefly, 1 × 10^6 cells per sample were stained in FACS buffer (BioLegend). Dead cells were excluded using a fixable viability dye (Invitrogen, L34979), and nonspecific antibody binding was blocked with mouse Fc Block (BD Biosciences, 553142). Cell subsets were identified according to the gating strategy shown in Supplementary Figure S2 and Supplementary Table S2. Briefly, B cell and T cell populations were identified by flow cytometry using the following key markers: total B cells (CD19+), transitional T1 (CD19+ CD23- CD21/35- IgM+), transitional T2 (CD19+CD23+ CD21/35+ IgM+), follicular (CD19+ CD93- CD23+ CD21/35+), marginal zone (CD19+ CD93- CD23- CD21/35+), germinal center (CD19+ GL7+ CD95+), plasmablasts (CD19+ CD138+ MHC II+), plasma cells (CD19- CD138+ MHC II-), CD4+ T cells (CD4+ TCRβ+), and Tfh (CXCR5+ PD-1+ ICOS+). Population frequencies were calculated as a percentage of viable CD45+ cells and, where indicated, normalized to the isotype control group. Data were analyzed in FlowJo version 10.10.0, and figures and statistical analyses were generated in GraphPad Prism. Normally distributed data were analyzed by one-way ANOVA with Dunnett’s multiple-comparisons test, whereas non-normally distributed data were analyzed using the Kruskal-Wallis test with Dunn’s multiple-comparisons test.

### Mouse Influenza Virus Challenge Model

Female C57BL/6 mice (6-8 weeks old) were purchased from Charles River Laboratories and randomized into six groups (n=10/group, except vehicle control, n=9). Mice received 4540, POV, or isotype control MOTA at 10 mg/kg by intraperitoneal injection twice weekly throughout the study. Vehicle and unvaccinated control groups received PBS at 10 mL/kg on the same schedule. A normal control group remained untreated and unmanipulated.

At weeks 4 and 7, mice in the 4540, POV, MOTA, and vehicle control groups were immunized intramuscularly with 5 µg of mouse-adapted, BEI-inactivated H1N1 A/California/07/2009 virus. At week 10, vaccinated groups and the unvaccinated control group were challenged intranasally with 300 × CCID_50_ of H1N1 A/California/07/2009 virus.

Animals were monitored daily for 21 days after challenge for clinical signs of infection and body weight loss. Mice were euthanized if body weight loss reached ≥30% of pre-challenge weight for 2 consecutive days. Survival was analyzed using Kaplan-Meier curves with between-group comparisons by the Mantel-Cox log-rank test.

All animal procedures were performed in accordance with the National Institutes of Health Guide for the Care and Use of Laboratory Animals and were approved by the Institutional Animal Care and Use Committee of Utah State University.

### KLH Mouse Vaccination Study

Male C57BL/6J mice (6 weeks old; 19-25 g) were purchased from The Jackson Laboratory (Bar Harbor, ME, USA) and randomized into 3 treatment groups (n=5/group). All mice were immunized intramuscularly with 250 µg KLH (Thermo Fisher Scientific, 77600) on days 0 and 42. Beginning on day 14, mice received 4540, POV, or isotype control antibody (MOTA) at 10 mg/kg by intraperitoneal injection twice weekly for the duration of the study. Blood was collected by submandibular puncture on days 14, 42, 49, and 56, and plasma was isolated for analysis.

Anti-KLH IgG and IgM levels were measured by ELISA using Life Diagnostics kits (KLHG-1 and KLHM-1) with modifications to the manufacturer’s protocol. Plates were coated with KLH at 0.4 µg/mL, and plasma samples were diluted to fall within the assay dynamic range. Bound antibody was detected using HRP-conjugated anti-mouse IgG or anti-mouse IgM secondary antibodies (Jackson ImmunoResearch, 115-035-071 and 115-035-075), followed by development with TMB Ultra substrate (Thermo Fisher Scientific, 34028). Results were interpolated from a standard curve and reported as KLH U/mL. Data were analyzed using Brown-Forsythe and Welch ANOVA with Dunnett’s T3 multiple-comparisons test or, for nonparametric data, the Kruskal-Wallis test with Dunn’s multiple-comparisons test. All animal procedures were performed in accordance with the National Institutes of Health Guide for the Care and Use of Laboratory Animals and were approved by the Institutional Animal Care and Use Committee of Mispro Biotech Services.

### Human B-cell BAFF/APRIL stimulation assay

Human B cells were isolated from cryopreserved peripheral blood mononuclear cells (PBMCs; STEMCELL Technologies, 70025) by negative selection using the EasySep Human B Cell Isolation Kit (STEMCELL Technologies, 17954). Purified B cells were seeded at 2 × 10^5 cells per well in 6-well plates in 2 mL RPMI 1640 medium supplemented with L-glutamine, 10% fetal calf serum, 10 mM HEPES (pH 7.4), 0.1 mM non-essential amino acids, and 1 mM sodium pyruvate. Cells were stimulated for 3 days with soluble recombinant human CD40 ligand (1 µg/mL; R&D Systems or PeproTech) and recombinant human IL-2 (2 ng/mL; R&D Systems).

After stimulation, cells were washed and reseeded at 1 × 10^5 cells per well in medium containing IL-2 (2 ng/mL) together with recombinant human BAFF (5 µg/mL; R&D Systems), recombinant human APRIL (500 ng/mL; R&D Systems), or both, as indicated. No-BAFF control wells were cultured in the absence of BAFF. Where indicated, cells were treated with XT1185, belimumab, povetacicept, or the isotype control motavizumab at 50 µg/mL. Cells were cultured for the indicated time periods at 37°C in a humidified incubator with 5% CO2.

Cell viability was assessed at the indicated time points using the CellTiter-Glo 2.0 Cell Viability Assay (Promega, G9242) according to the manufacturer’s instructions. For luminescence measurement, CellTiter-Glo reagent was added directly to the cultures, and duplicate aliquots from each well were analyzed as technical replicates. Experiments were performed using cells from a single donor, with duplicate culture wells per condition

### Single cell CITE-seq data generation

Previously dissociated and cryopreserved mouse splenocytes treated with APRIL inhibitor (4540), APRIL/BAFF inhibitors (POV or ATA), or isotype control antibody (MOTA) were processed following 10x sample preparation guidelines. Briefly, thawed cells were stained with Trustain FcX anti-mouse CD16/32 antibody (Biolegend Cat # 101320) for 5 min and incubated with TotalSeq-C anti-mouse hashtags and washed, pooled in equal numbers from each condition, and stained with Biolegend’s Totalseq C Universal cocktail (Cat #199903) plus spike-in antibodies , resulting in a total of 129 proteins targeted for detection (Supplementary Table S3 and S4). Four pools from 17 samples were generated, resulting in 14 libraries being prepared using 10x GEM-X single cell 5’ V3 chemistry to target 10,000 cells per sample (Supplementary Table S5). The quality of final libraries was assessed via Bio Agilent Fragment Analyzer using 1-6000bp NGS standard sensitivity kit and sequenced on Illumina NovaSeq 2x150bp configuration, targeting for 30,000 and 10,000 read pairs per cell for gene expression and feature barcode libraries, respectively.

### Xenium spatial transcriptomics data generation

To profile the spatial organization of gene expression at subcellular resolution, we utilized the Xenium Analyzer In Situ platform (10x Genomics, Pleasanton, CA) following the manufacturer’s protocol. Briefly, mouse spleens for spatial analysis were fixed for 144 hours in 10% Neutral Buffered Formalin at 4-degrees. Following fixation, tissues were processed through a standard ethanol dehydration series, cleared with xylene, and embedded in paraffin blocks. Sections of 5 µm thickness were cut and mounted onto Xenium slides, followed by deparaffinization, enzymatic digestion, and hybridization with the Xenium Prime 5K Mouse Pan Tissue and Pathways Panel (10X Genomics, PN-1000725). The bound probes were subsequently amplified via rolling circle amplification. Slides were then loaded onto the Xenium Analyzer for automated cycles of fluorescent secondary probe hybridization, high-resolution imaging, and decoding chemistry. Sections were stained with H&E upon run completion.

### Single cell CITE-seq data analysis

Single-cell 5′ gene expression data were processed with Cell Ranger (v7.0) against the mouse reference GRCm39-2024-A, and downstream analyses were performed in R (v4.3.3) using Seurat (v5)^80^. After batch-wise quality control (100 < nFeature_RNA < 6,000; percent.mt < 15%) and hashtag-based demultiplexing with an adapted HTODemux workflow, datasets were merged, library adjusted and log normalized and scaled. A variance stabilizing transformation was used to identify the top 2,000 highly variable genes before applying principal components analysis (PCA) with the top 30 PCs used for downstream analyses. Doublets were identified with DoubletFinder (v2.0.6)^81^, batch effects were corrected using Harmony^82^ on the first 30 PCs, and UMAP embeddings were computed from the integrated space. An additional mitochondrial filter (percent.mt < 5%) was applied following inspection, yielding 141,811 cells for analysis. Cell identities were assigned using canonical markers informed by both RNA and ADT measurements. Differential gene expression analyses were performed in a pseudobulk framework by aggregating counts per sample and cell subtype via DESeq2 (v1.42.0)^83^, incorporating a log(nCells) offset for dual APRIL/BAFF inhibition samples to account for treatment-induced B-cell depletion. Cell abundances were calculated relative to total T cells and compared between groups using two-sided Mann–Whitney U tests. Fina figures were generated in R with ggplot2 (v4.0) and Seurat. See Supplementary Methods for additional information.

### Xenium data analysis

Spatial transcriptomics data were generated using the 10x Genomics Xenium platform, with imaging, barcode decoding, and cell segmentation performed by Xenium Ranger (v3.3.0.1). Count matrices from all samples were merged into a single Anndata object (Scanpy v1.11.5)^84^ comprising of > 4 million cells; downstream analyses and spatial data handling were performed using SpatialData (v0.4.0)^85^. Cells with fewer than 70 detected genes were excluded. For dimensionality reduction and clustering, a balanced working subset of 25,000 cells per sample was generated to maintain representation across treatment conditions. Counts were library-size adjusted and log-normalized. PCA was performed using default settings, and k-nearest neighbor graphs (k = 15) were constructed for UMAP embedding and Leiden clustering. Clustering resolutions were performed across resolutions from 0.1 to 1.5 in increments of 0.1 to identify an optimal resolution that separated known immune lineages without over-clustering. Cell type annotation was guided by canonical lineage markers and differential expression analysis (see Supplementary Methods). Annotations were transferred to the full dataset using Scanpy’s k-nearest neighbor label transfer procedure (sc.tl.ingest), and cell-type proportions between the annotated subset and full dataset were verified to be consistent. Cell-type proportions per sample were calculated as the fraction of cells assigned to each type relative to total detected cells. Germinal centers were identified within B-cell zones by computing spatial kernel density estimates of *Aicda* and *Mki67* expression intersected with germinal center B cell annotations; candidate regions were then manually verified and quantified across samples and treatment conditions. See Supplementary Methods for additional information.

## SUPPLEMENTAL MATERIAL CONTENTS

### Supplemental methods

Extended single cell CITE-seq data analysis

Extended Xenium spatial transcriptomics data analysis

Flow cytometry analysis of B-cell subsets from Sibeprenlimab ENVISION trial

Supplemental Figures S1-S11

Figure S1. Mouse bodyweight trajectory post influenza viral challenge.

Figure S2. Impact of APRIL and APRIL/BAFF inhibition on serum IgG and IgM after 8 weeks of treatment.

Figure S3. Flow cytometry gating schemes for spleen immunophenotyping. Figure S4. Supporting marker genes of non-B-cell populations.

Figure S5. RNA and ADT expression show agreement in defining B-cell subpopulations.

Figure S6. Spatial transcriptomics data quality and annotation.

Figure S7. Mixed cell annotations exist at non-random cell interfaces, representing distinct microarchitectures of the spleen.

Figure S8. Signaling proteins and genes between follicular B and T cells are modulated by APRIL and APRIL/BAFF inhibition.

Figure S9. Extended B cell-T cell costimulation and chemotaxis genes

Figure S10. Differential gene expression analysis reveals minimal differences between GC cells in 4540 vs isotype.

Figure S11. APRIL inhibition spares class-switched memory B cells in human clinical trial.

## Supplemental Tables S1-S5

Table S1: Antibodies and reagents used for flow cytometric immunophenotyping. Table S2: Flow cytometry gating definitions for immune cell populations.

Table S3: ADTs in Biolegend’s Totalseq C Universal cocktail excluding isotype controls. Table S4. Additional Spike-in CITE-seq Antibodies.

Table S5. Mouse splenocyte sample pools for single cell CITE-seq analysis.

## ACKNOWLEDGEMENTS

The authors thank the Visterra In Vivo Biology team for supporting the mouse studies and the Institute for Viral Research at Utah State University for support of the influenza challenge model. The authors also thank Zach Shriver for scientific perspective and discussion of the work and manuscript, and Prashanna Balaji Venkatasubramanian for assistance with single-cell analyses.

The authors thank the patients and investigators who participated in the ENVISION trial. This work was funded by Visterra, Inc.

## DATA SHARING STATEMENT

Data will be shared upon publication in a peer-reviewed journal.

## AUTHOR CONTRIBUTIONS

KR, GJB, and LNR provided substantial contributions to the conception of the work. KR, VS, AR, KAH, JB, JVK, DF, ML, BH, and LNR substantially contributed to the acquisition, analysis, or interpretation of data for the manuscript. KR, GJB and LNR drafted the manuscript; all authors critically reviewed and approved final version of this manuscript to be published and accept accountability for the work.

## USE OF ARTIFICIAL INTELLIGENCE

Artificial intelligence tools were used to assist with editing for clarity and language refinement during manuscript preparation. Scientific content, analysis, and interpretation were performed by the authors.

