## Supplementary Material for "Targeting the APRIL-BAFF axis in IgA nephropathy: preclinical insights from influenza challenge and single-cell spatial profiling"

### SUPPLEMENTAL METHODS

#### Extended single cell CITE-seq data analysis

*Single-cell RNA-seq preprocessing*

10x Genomics Single Cell 5′ gene expression data were processed with Cell Ranger (v7.0) against the mouse reference GRCm39-2024-A provided by 10x Genomics. Libraries were generated across four batches. For each batch, we retained barcodes with 100 < nFeature_RNA < 6,000 and percent mitochondrial transcripts (percent.mt) < 15%, then performed sample demultiplexing using an in-house adaptation of Seurat's HTODemux with CLR-normalized ADTs, mitigating hashtag spillover during classification. The resulting batch-level objects were merged, yielding 155,878 cells prior to doublet removal.

Downstream analyses were conducted in R (v4.3.3) using Seurat (v5)[1]. RNA data were normalized with NormalizeData and scaled with ScaleData. We identified 2,000 highly variable genes and computed principal component analysis (PCA) on these features; the number of informative PCs was selected using the elbow plot and cumulative variance explained, and the top 30 PCs were carried forward. Putative doublets were identified and removed with DoubletFinder (v2.0.6)[2] using default parameters. Batch effects were mitigated with Harmony (v1.0)[3] on the first 30 PCs (max.iter.harmony = 30; other parameters default), and UMAP was computed from the same 30-dimensional embedding using Seurat defaults. Following doublet removal, we observed small islands with elevated mitochondrial content and therefore applied a more stringent QC step to exclude cells with percent.mt > 5%, consistent with common mouse scRNA-seq practice and data-driven inspection of our dataset (reference note below). Cell subsets were annotated using canonical marker genes, leveraging both RNA and ADT measurements to refine labels. After QC, demultiplexing, doublet removal, integration, and filtering, 141,811 cells remained for downstream analyses.

*Differential expression analysis*

Differential expression was performed in a pseudobulk framework. For each cell subtype within each sample (aggregation unit = sample × cell subtype), raw counts were summed to form pseudobulk profiles and analyzed with DESeq2 (v1.42.0)[4]. Genes were retained if they had at least 10 counts in at least two pseudobulk samples. To account for treatment-induced B-cell ablation in the dual APRIL/BAFF inhibition arm, we included a log(nCells) offset only for those pseudobulk samples, leaving the offset as zero for other arms. DESeq2 was otherwise run with default settings using sfType = "poscounts"; log fold-change shrinkage was applied with lfcShrink under default parameters; and multiple testing correction followed the DESeq2 defaults.

*Abundance analysis*

Cell abundances were estimated by normalizing B-cell subsets, Tfh, and Tfr against total T cells. This denominator was chosen a priori because T-cell numbers were not reduced by treatment, whereas B cells were heavily depleted under dual inhibition, allowing T cells to serve as a relatively stable reference population. Group differences between treatments and controls were assessed at the sample level using two-sided, non-parametric Mann–Whitney U tests in R.

*Figure generation*

Figures were generated in R using ggplot2 (v4.0.0); UMAP visualizations were produced with Seurat in conjunction with RColorBrewer (v1.1.3) for palettes.

**Extended Xenium data analysis**

*QC and pre-processing*

Imaging, barcode decoding, and cell segmentation were performed with Xenium Ranger (v3.3.0.1). Summary reports generated by Xenium Ranger confirmed high quality imaging and decoding across all runs, with no runtime errors and appropriate cell segmentation.

Distributions of detected genes and transcripts per cell were comparable across samples and treatment arms. These distributions informed quality control thresholds used during initial processing. The ≥70-gene cutoff was selected to remove lower-information cells that contributed to noisy or poorly defined clusters. We applied this conservative gene-content filter along with sub-sampling to annotate cell types, in which the labels were propagated the remaining cells. We verified that the proportions of each annotated cell type were balanced both in the subset data and the label transferred result.

*Dimensionality reduction and clustering*

Count matrices from all samples were merged into a single Anndata (Scanpy v1.11.5)[5] object covering more than 4 million cells. For dimension reduction and clustering, we created a balanced working subset by randomly sampling 25,000 cells per sample. This subset-maintained representation across treatment conditions while enabling more efficient computation. PCA and UMAP embeddings suggested no observable batch effects and therefore no batch correction was applied to the embeddings. Leiden clustering was performed at increasing resolutions from 0.1 to 1.5 at increments of 0.1 to define an optimal resolution that separated known immune lineages and subtypes while avoiding over clustering. Cluster labels projected back into tissue coordinates showed spatial patterns consistent with spleen anatomy and did not identify biases introduced by the analysis.

*Cell type annotation and label transfer*

Annotation of the down-sampled dataset set was led by canonical lineage markers and DE analysis, using Scanpy's *rank_genes_groups* function. High-level lineages included B cells (*Cd79a*, *Ms4a1*, *Cd19*), T cells (*Cd8b1*, *Trac*, *Cd3e*), myeloid (*Clec9a*, *Siglec1*, *Cd14*, *Cd68*), and stromal cells (*Cxcl12*, *Col1a2*). B cells could further be split into with distinct localization in tissue, including antibody secreting cells and germinal center B cells. Mixed populations were observed at structural interfaces, where co-expression of expected non-overlapping lineage markers (e.g., *Cd79a* and *Cd3e*) were found and therefore annotated as mixed-lineage clusters for their ambiguity. Labels were transferred back to the full dataset using sc.tl.ingest, which performs k-nearest neighbor mapping in principal component embeddings. Comparing cell-type proportions between the annotated working set and the label-transferred full set revealed no immediate bias of these results.

*Proportion analyses*

Cell-type proportions were estimated on the full, label-transferred dataset as the number of cells assigned to each cell type divided by the total number of detected cells in that sample. These per-sample proportions were used to compare treatment-wise differences in cellular composition.

*Spatial expression patterns and germinal center quantification*

Germinal center activity was defined as regions within B-cell zones where localized expression of *Mki67* and *Aicda* were identified. To identify these in tissue, we computed spatial kernel density estimates of *Aicda* and identified the intersect with germinal center B-cell annotations. The combination of these features facilitated quicker manual annotation of existing germinal centers in each sample. Two-sided exact permutation tests were used to test for significant changes in germinal center counts under April or April/Baff dual inhibition compared to PBS and isotype controls.

*Cell-cell colocalization analyses*

Using spatial cell coordinates from a representative 4540-treated sample, we performed a nearest-neighbor analysis to identify the 10 closest neighboring cells for each cell in tissue. Neighbor relationships were summarized by cell type, then row-normalized to characterize local cell adjacency and visualized as a heatmap.

*Signal contamination assessment in spatial transcriptomics data*

A recognized technical artifact of in situ hybridization (ISH)-based spatial transcriptomics platforms is the potential for transcript mis-assignment between adjacent cells, arising from transcript diffusion between closely apposed cells and/or errors in cell segmentation[6, 7]. In our Xenium spatial transcriptomics dataset, we identified a cluster that exhibited co-expression of canonical B cell markers (*Cd79b*, *Ms4a1*) alongside T cell markers (*Cd3e*). In UMAP space, this cluster was positioned between the major B cell and T cell clusters, suggesting intermediate gene expression characteristics. Inspection of the spatial localization of cells assigned to this cluster revealed that they were concentrated at the boundaries between B cell-rich and T cell-rich regions of follicles, precisely where transcript diffusion between neighboring B and T cells would be expected to be most prevalent. These observations are consistent with the signal contamination patterns reported by others using ISH-based spatial transcriptomics technologies, in which enrichment of mis-assigned transcripts occurs preferentially among cells in close physical proximity, potentially due to transcript diffusion across cell membranes or imprecise computational cell segmentation. The consistent localization of this mixed-marker cluster at B-T cell boundaries, rather than in random tissue locations, further supports a technical rather than biological origin for the mixed expression pattern. Our clustering identified these mixed populations, which were annotated as “mix” (Supplemental Figure 7A).

**Flow Cytometry Analysis of B Cell Subsets from Sibeprenlimab ENVISION trial**

Peripheral blood B-cell subsets were analyzed by flow cytometry in a subset of participants from the ENVISION trial (NCT04287985), a phase 2, randomized, placebo-controlled study of intravenous sibeprenlimab (2, 4, or 8 mg/kg monthly) in adults with IgA nephropathy. Full study design, eligibility criteria, clinical results, and analyses of total B-cell and T-cell populations are reported in (McCafferty et al, manuscript under review). The trial was conducted in accordance with the principles of the Declaration of Helsinki and the International Council for Harmonisation Guideline for Good Clinical Practice. The institutional review board or ethics committee for each participating center approved the protocol before initiation of the trial. The isotype-resolved memory B-cell subset analyses presented here (IgA+, IgM+, and inferred IgG+ memory B cells) have not been previously reported. Samples were collected on Day 1 (baseline), Day 180, Day 360 (end of treatment), and Day 485 (follow-up).

Whole blood samples were stained with a B-cell class switch panel comprising antibodies against CD3, CD14, CD19, CD27, CD38, CD43, IgA, IgD, and IgM, and acquired on a BD FACSCanto II flow cytometer. CD38 and CD43 were included in the panel for broader B-cell subset characterization but were not used in the gating strategy for the analyses presented here. Following exclusion of debris and doublets, lymphocytes were gated by forward and side scatter. CD3+ T cells and CD14+ monocytes were excluded, and CD19+ B cells were identified from the remaining population. Memory B cell subsets were analyzed (CD19+ CD27+), and IgA+ and IgM+ memory B cells were identified by surface immunoglobulin staining. IgG+ memory B cells were inferred by exclusion of IgA and IgM surface staining from the memory B-cell gate (CD19+CD27+IgM−IgA−), as the panel did not include a direct anti-IgG reagent (a minor fraction of IgA−IgM− class-switched memory B cells may include IgE+ cells, though these are typically rare in peripheral blood).

### SUPPLEMENTAL FIGURES

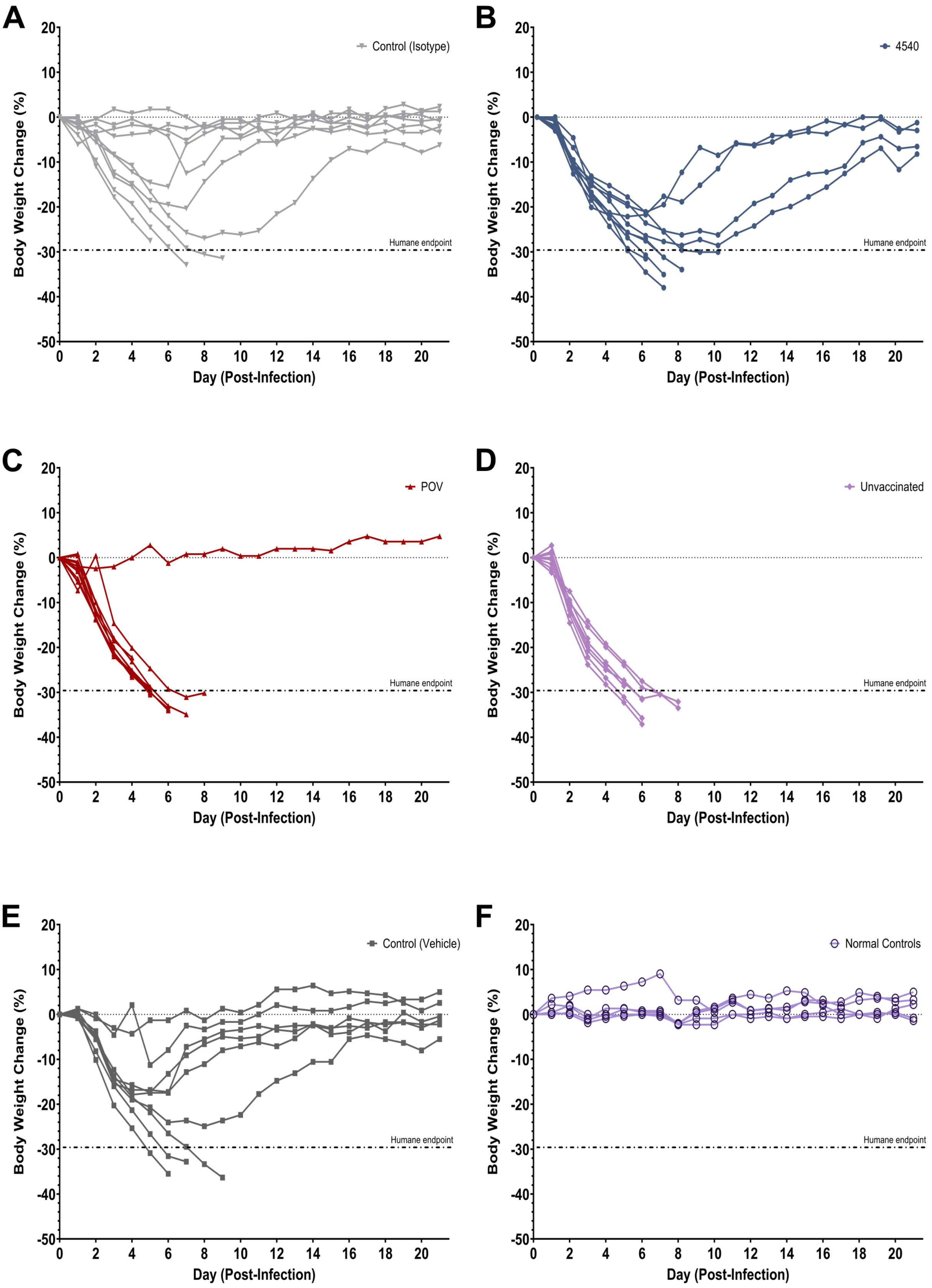
A
Control (Isotype)
Humane endpoint
Body Weight Change (%)
Day (Post-Infection)
B
4540
Humane endpoint
Body Weight Change (%)
Day (Post-Infection)
C
POV
Humane endpoint
Body Weight Change (%)
Day (Post-Infection)
D
Unvaccinated
Humane endpoint
Body Weight Change (%)
Day (Post-Infection)
E
Control (Vehicle)
Humane endpoint
Body Weight Change (%)
Day (Post-Infection)
F
Normal Controls
Humane endpoint
Body Weight Change (%)
Day (Post-Infection)

**Figure S1. Mouse bodyweight trajectory post influenza viral challenge.**

Individual body-weight trajectories are shown for mice in the **A)** isotype control, **B)** APRIL inhibitor (4540), **C)** APRIL/BAFF dual inhibitor (POV), **D)** unvaccinated, **E)** vehicle control, and **F)** healthy control groups. Body weight was recorded daily from day 0 (viral challenge) through day 21 post-infection and is presented as percent change from each animal's day 0 body weight. Each line represents one animal. The dashed line denotes the humane endpoint threshold of 30% body-weight loss. Animals with >30% loss of baseline body weight for two consecutive days were humanely euthanized and removed from the study

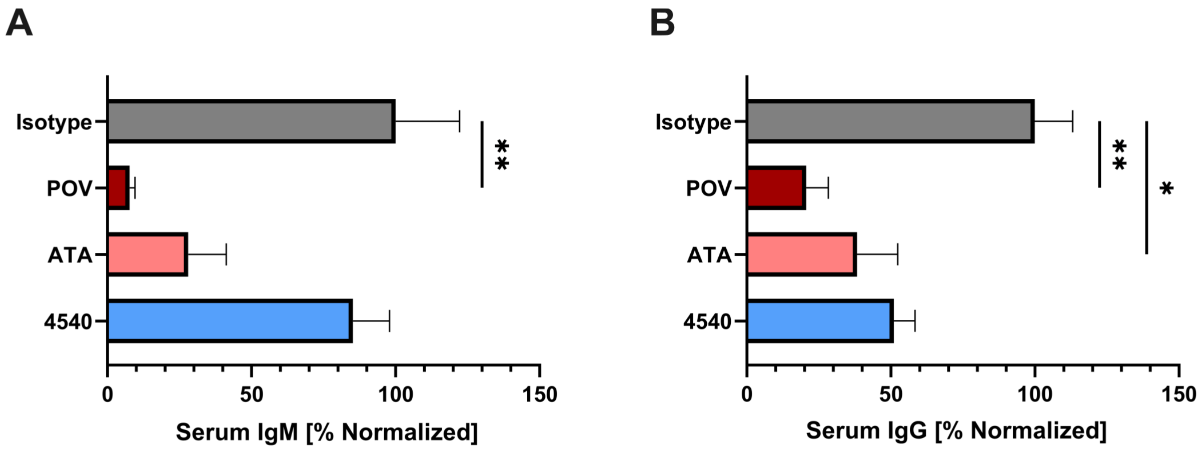


**Figure S2. Impact of APRIL and APRIL/BAFF inhibition on serum IgG and IgM after 8 weeks of treatment.**

Serum **A)** IgM, and **B)** IgG levels measured at week 8 in mice treated with APRIL inhibitor (4540), APRIL/BAFF dual inhibitors (POV or ATA), or isotype control. Titers are normalized relative to the average titer of the isotype control group. Bars represent mean ± SEM. Statistical analysis was performed using Kruskal-Wallis test with Dunn's multiple-comparisons test *p = <0.05, **p = <0.01.

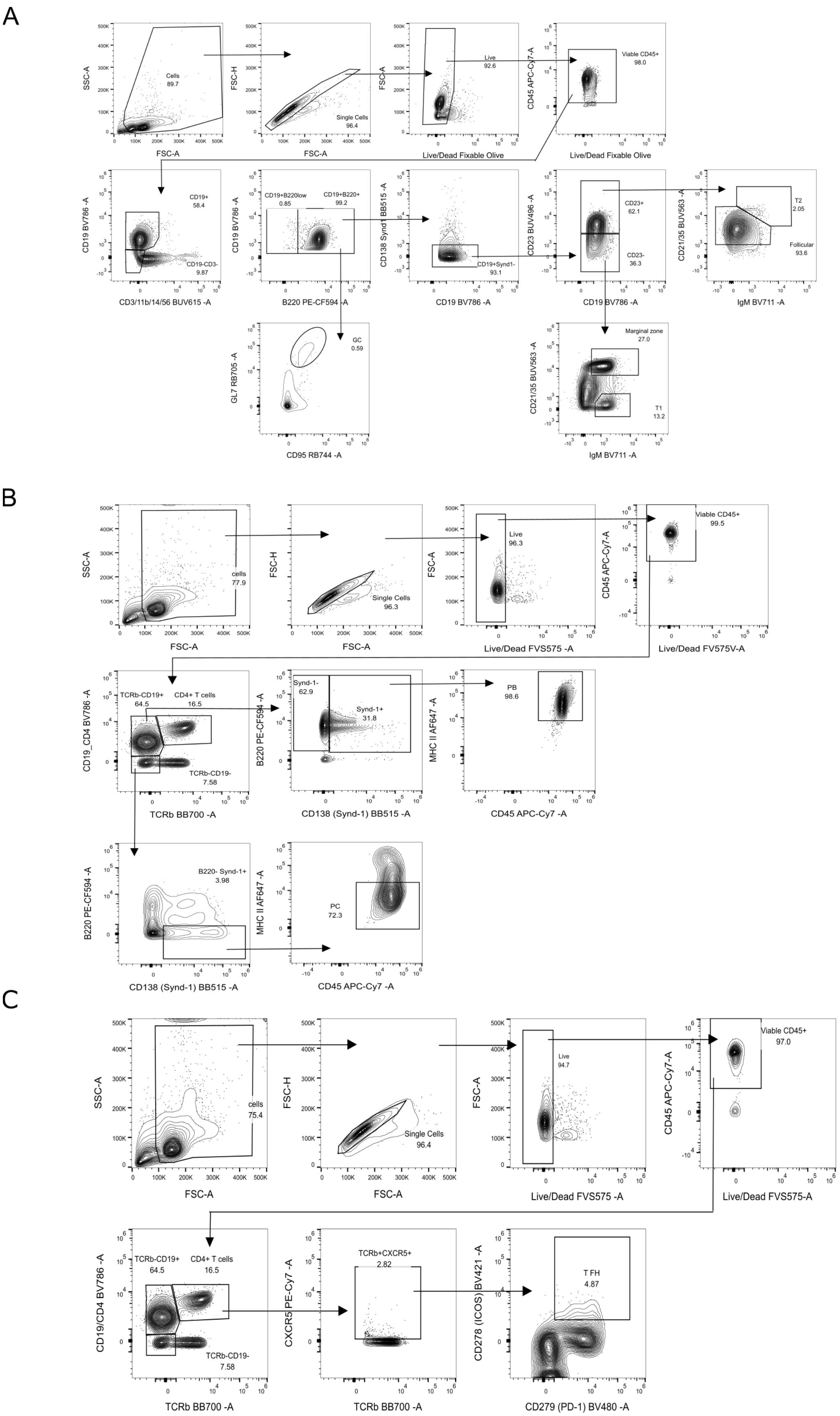
A
Cells 89.7
SSC-A
FSC-A
FSC-H
Single Cells 96.4
Live 92.6
Live/Dead Fixable Olive
CD45 APC-Cy7-A
Viable CD45+ 98.0
CD19 BV786 -A
CD19+ 58.4
CD19-CD3- 9.87
CD3/11b/14/56 BUV615 -A
CD19+B220low 0.85
CD19+B220+ 99.2
B220 PE-CF594 -A
CD138 Synd1 BB515 -A
CD19+Synd1- 93.1
CD23 BUV496 -A
CD23+ 62.1
CD23- 36.3
CD21/35 BUV563 -A
T2 2.05
Follicular 93.6
IgM BV711 -A
GL7 RB705 -A
GC 0.59
CD95 RB744 -A
Marginal zone 27.0
T1 13.2
B
cells 77.9
Single Cells 96.3
Live 96.3
Live/Dead FVS575 -A
Viable CD45+ 99.5
Live/Dead FV575V-A
CD19_CD4 BV786 -A
TCRb-CD19+ 64.5
CD4+ T cells 16.5
TCRb-CD19- 7.58
TCRb BB700 -A
Synd-1- 62.9
Synd-1+ 31.8
CD138 (Synd-1) BB515 -A
MHC II AF647 -A
PB 98.6
CD45 APC-Cy7 -A
B220- Synd-1+ 3.98
PC 72.3
C
cells 75.4
Single Cells 96.4
Live 94.7
Live/Dead FVS575 -A
Viable CD45+ 97.0
Live/Dead FVS575-A
CD19/CD4 BV786 -A
TCRb+CXCR5+ 2.82
CXCR5 PE-Cy7 -A
CD278 (ICOS) BV421 -A
T FH 4.87
CD279 (PD-1) BV480 -A

**Figure S3. Flow cytometry gating schemes for spleen immunophenotyping.**

Individual populations of **A)** B cells, **B)** Antibody-secreting and **C)** T cell subsets. Gates were set utilizing fluorescent minus one (FMO) conditions. Boxes denote the percentage of cells as a subset of the parental gate.

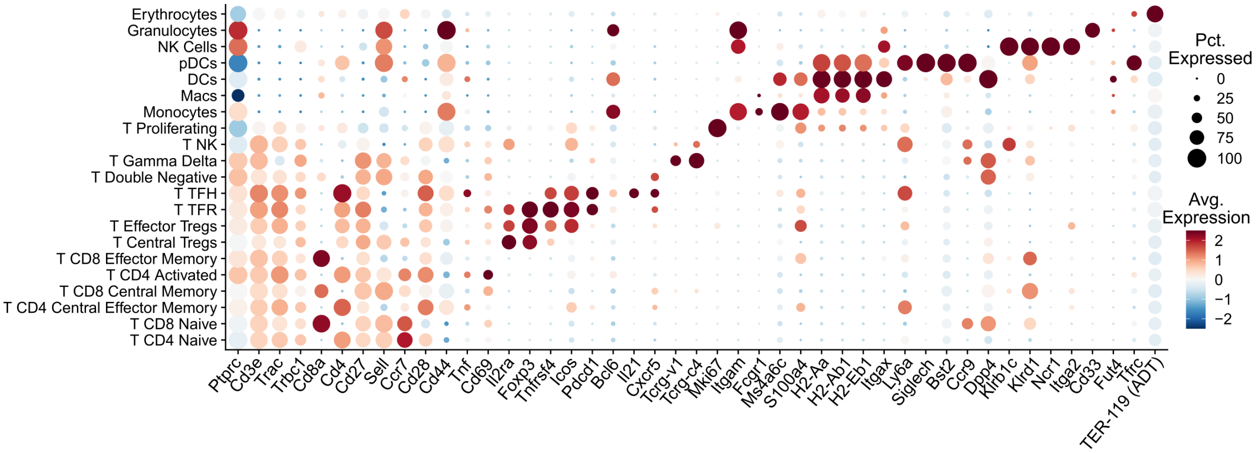


**Figure S4. Supporting marker genes of non-B cell populations.**

Dot plot showing expression of selected marker genes across non-B cell clusters including T cells, myeloid cells, and others. Color indicates normalized average gene expression and size reflects the percentage of cells within each cluster expressing the marker. TER-119 ADT expression was used to highlight erythrocytes since gene expression was not captured.

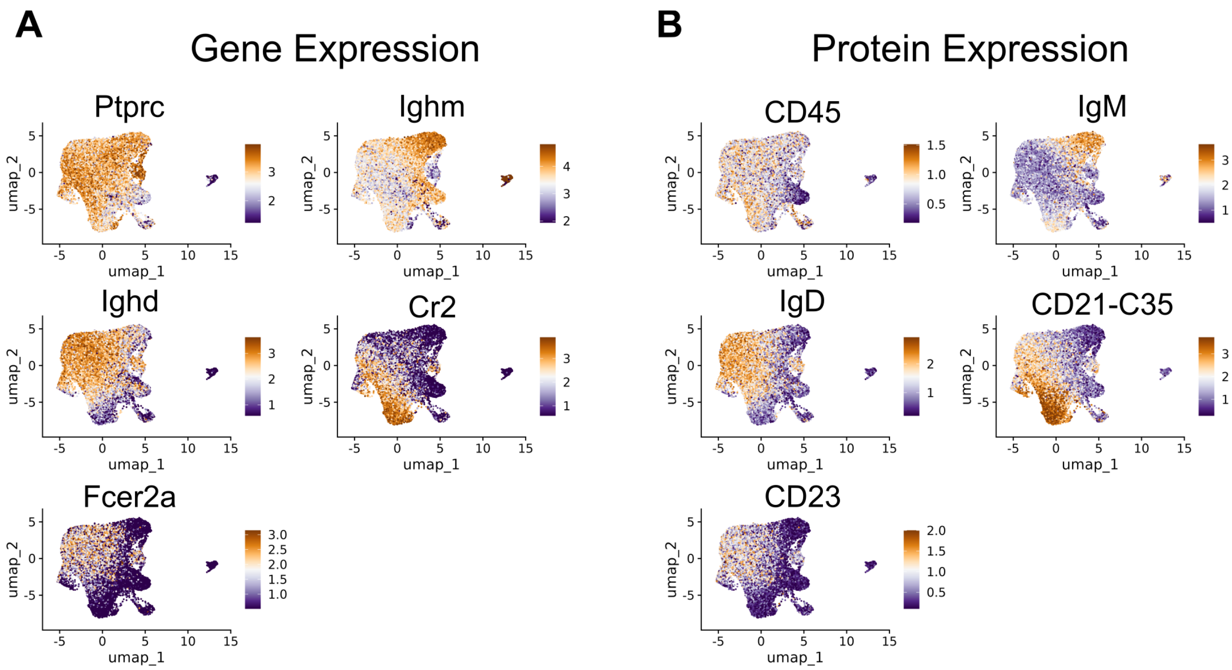


**Figure S5. RNA and ADT expression show agreement in defining B cell subpopulations.**

B cell UMAPs showing **A)** gene expression and matched **B)** protein (ADT) expression for select B cell markers.

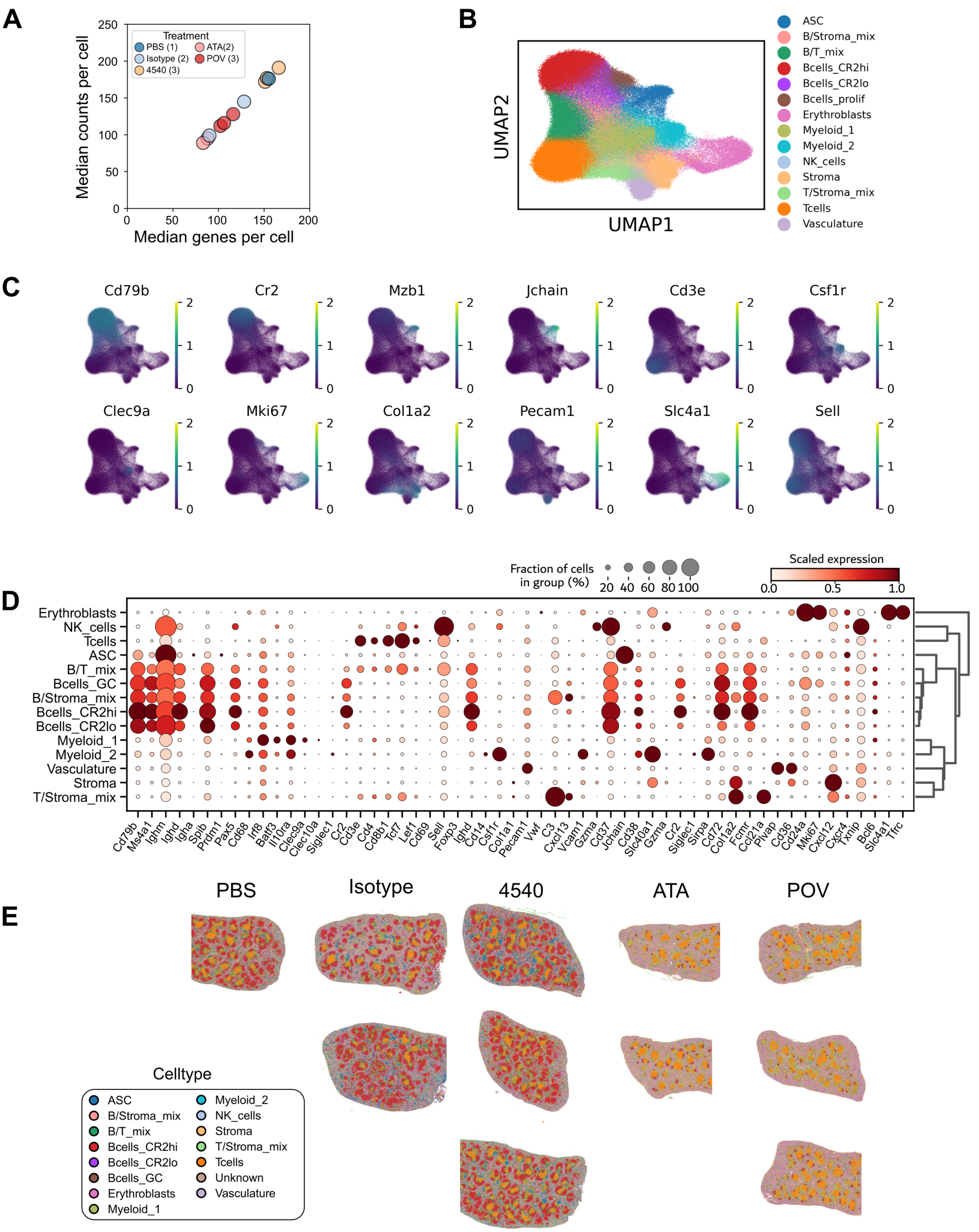


195

196 **Figure S6. Spatial transcriptomics data quality and annotation.**

**A)** Per sample quantification of median transcript and gene counts from Xenium transcriptomics analyses. **B)** UMAP embeddings and cell clusters were generated on a merged dataset of control and treated spleens (1 PBS, 2 MOTA, 3 4540, 2 POV, and 3 ATA). Each spleen was randomly down sampled to 25k cells for visualization, and computational efficiency and annotated using lineage specific genes. **C)** Feature plots highlight subset specific cells for B cell phenotypes (CD79b, Cr2, Mzb1, Jchain), pan T cells (Cd3e), myeloid cells (Csf1r, Clec9a) proliferation (Mki67), stroma and vasculature (Col1a2, Pecam1), and immune trafficking (Sell). Values represent log-normalized expression. **D)** Visualization of selected genes from lineage markers and differential expression analyses. The color intensity represents per marker scaled expression ranges from 0-1. **E)** Cell type annotations were labeled in each sample showing consistency within treatment groups.

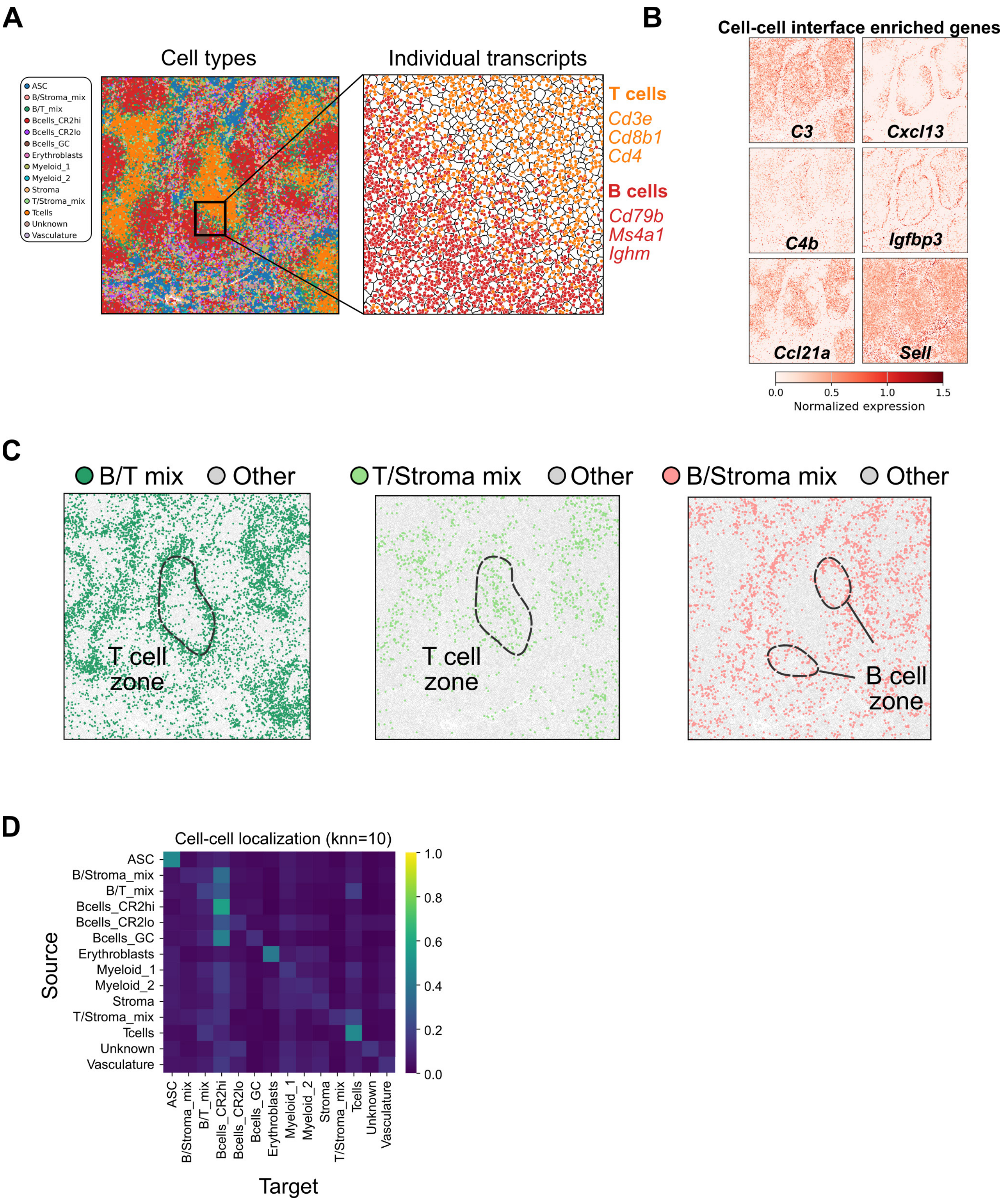


**Figure S7. Mixed cell annotations exist at non-random cell interfaces, representing distinct microarchitectures of the spleen.**

**A)** Cell type annotations derived from clustering overlaid onto corresponding spatial coordinates in a representative control spleen. Individual transcripts were visualized at the interface of B and T cell zones to show mixing of lineage-specific transcripts. **B)** Cells at B/T and stroma/immune interfaces are enriched for a subset of genes, representing functionally relevant regions in tissue. Color represents mean log normalized expression. **C)** Mixed cell type populations were visualized independently to highlight their non-random spatial organization. **D)** The top 10 spatial nearest neighbors for each cell type were averaged to show a seed cell type’s probability of being a neighbor to a target cell type. Results show row-normalized values (sum to 1).

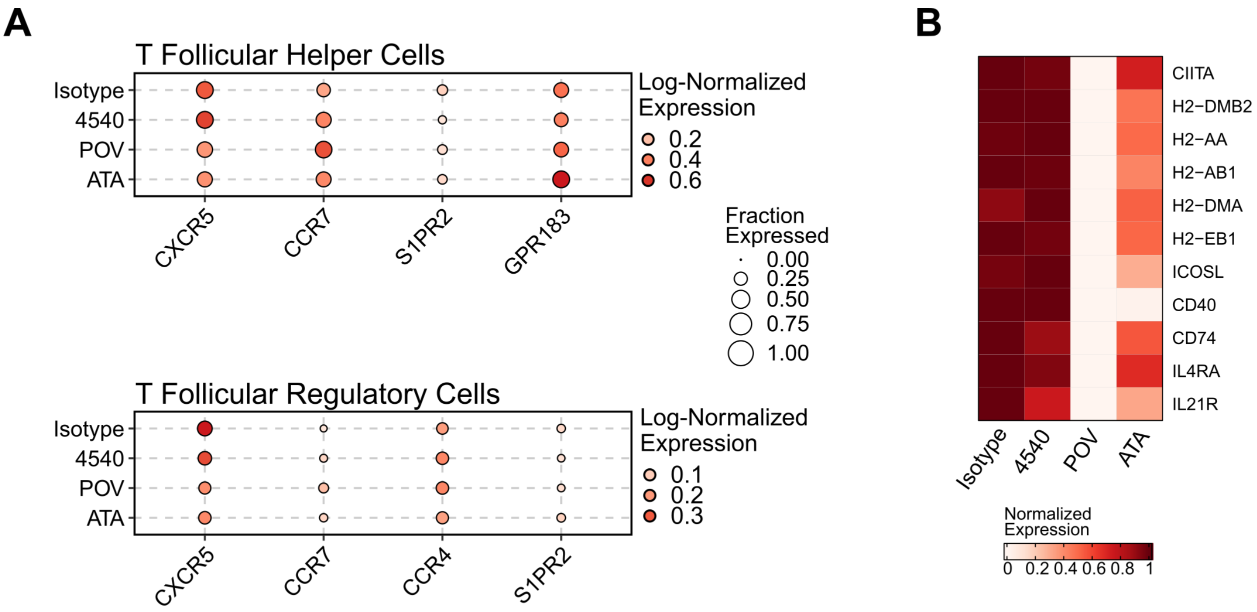


**Figure S8. Signaling proteins and genes between follicular B and T cells are modulated by APRIL and APRIL/BAFF inhibition.**

**A)** Average log-normalized expression in Tfh and Tfr cells for chemokine and trafficking receptors involved in GC formation across treatment groups. **B)** Heatmap of key genes mediating B cell–T follicular helper (TFH) cell interactions in follicular (FO) B cells. Values are scaled from 0 to 1.

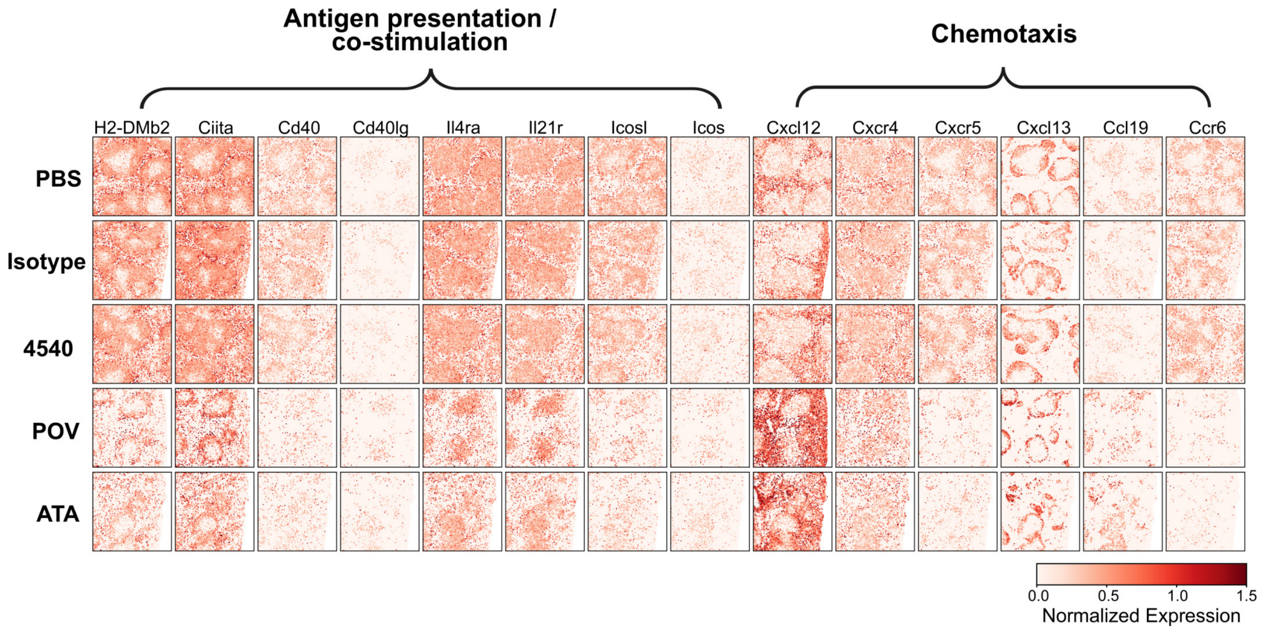


**Figure S9. Extended B cell-T cell costimulation and chemotaxis genes**

Extended list of relevant costimulation (B-T cell interaction genes) and chemotaxis genes represented in spleen tissue. Values are log-normalized expression.

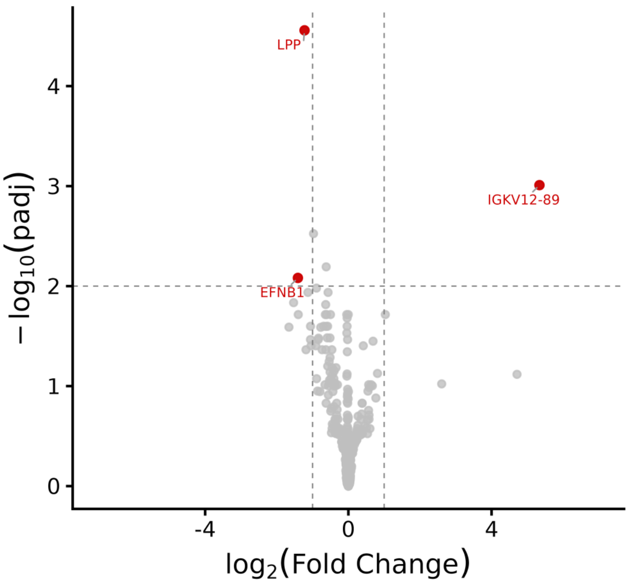


**Figure S10. Differential gene expression analysis reveals minimal differences between GC cells in 4540 vs isotype.**

Volcano plot of differentially expressed genes from single-cell pseudobulked GC B cells. Upregulated genes are increased in 4540 compared to isotype, fold change and p-values were calculated using DESeq2's Wald test.

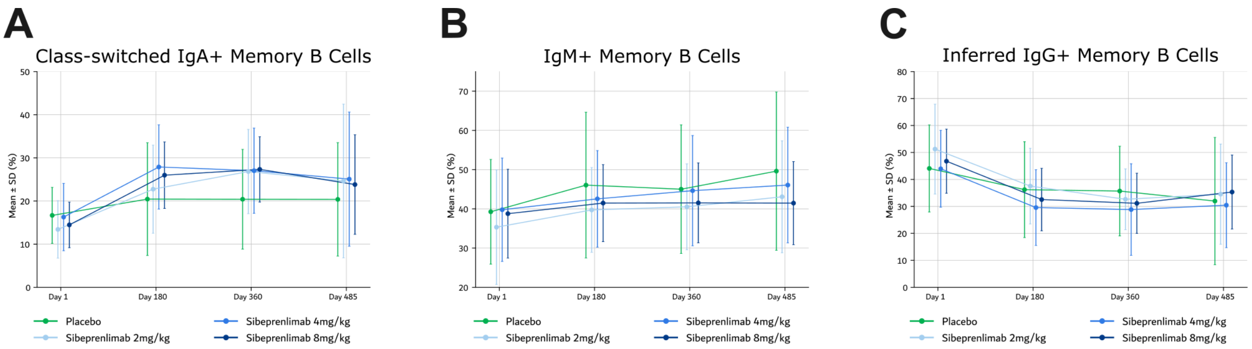


**Figure S11. APRIL inhibition spares class-switched memory B cells in sibeprenlimab-treated patients with IgA nephropathy.**

Peripheral blood memory B-cell subsets in sibeprenlimab-treated and placebo-treated patients with IgA nephropathy from the ENVISION trial (NCT04287985). **A)** IgA+ memory B cells (CD19+CD27+IgA+), **B)** IgM+ memory B cells (CD19+CD27+IgM+), and **C)** inferred IgG+ memory B cells (CD19+CD27+IgM−IgA−) were quantified by flow cytometry over 12 months of treatment and a subsequent follow-up period. Full gating strategy and flow cytometry methods are described in Supplementary Methods. The flow cytometry panel and ENVISION trial design are detailed in [McCafferty et al, manuscript under review].

### SUPPLEMENTAL TABLES

**Table S1: Antibodies and reagents used for flow cytometric immunophenotyping**

| **Marker** | **Color/Format** | **Catalog #** | **Clone** | **Titration** | **Manufacturer** |
|---|---|---|---|---|---|
| CD56 | Brilliant Ultraviolet 615 | 751046 | 809220 | 1:100 | BD Biosciences |
| CD3 | Brilliant Ultraviolet 615 | 751443 | 145-2C11 | 1:100 | BD Biosciences |
| CD4 | BV786 | 563727 | RM4-5 | 1:400 | BD Biosciences |
| CD11b | Brilliant Ultraviolet 615 | 751140 | M1/70 | 1:100 | BD Biosciences |
| IgM | Brilliant Violet 711 | 747732 | R6-60.2 | 1:50 | BD Biosciences |
| CD19 | Brilliant Violet 786 | 563333 | 1D3 | 1:400 | BD Biosciences |
| CD138 | Brilliant Blue 515 | 564511 | 281-2 | 1:100 | BD Biosciences |
| CD45R (B220) | PE-CF594 | 562290 | RA3-6B2 | 1:800 | BD Biosciences |
| MHC II | AlexaFluor 647 | 562367 | M5/114.15.2 | 1:800 | BD Biosciences |
| CD45 | APC-Cy7 | 557659 | 30-F11 | 1:100 | BD Biosciences |
| CD95 | RealBlue 744 | 756863 | Jo2 | 1:100 | BD Biosciences |
| CD23 | Brilliant Ultraviolet 496 | 741058 | B3B4 | 1:100 | BD Biosciences |
| GL7 | RealBlue 705 | 570602 | GL7 | 1:200 | BD Biosciences |
| CD21/CD35 | Brilliant Ultraviolet 563 | 752914 | 7E9 | 1:100 | BD Biosciences |
| CD14 | Brilliant Ultraviolet 615 | 756783 | SA14-2 | 1:100 | BD Biosciences |
| TCRb | BB700 | 745846 | H57-597 | 1:200 | BD Biosciences |
| CXCR5 | PE-Cy7 | 560617 | 2G8 | 1:50 | BD Biosciences |
| CD278 (ICOS) | BV421 | 564070 | 7E.17G9 | 1:200 | BD Biosciences |
| CD279 (PD1) | BV480 | 746784 | J43 | 1:100 | BD Biosciences |
| Live/Dead Fix Olive | Live/Dead Fix Olive | L34979A | --- | 1:1000 | Invitrogen |

| | | | | | |
|---|---|---|---|---|---|
| Fc Block | --- | 553142 | 2.4G2 | 1:1000 | BioLegend |

**Table S2: Flow cytometry gating definitions for immune cell populations**

| **Cell population** | **Cell Markers** |
|---|---|
| Total B cell | CD45+ CD19+ CD3- CD11b- CD14- CD56- |
| T1 | CD45+ CD19+ CD3- CD11b- CD14- CD56- B220+ CD138- CD23- CD21/35- IgM+ |
| T2 | CD45+ CD19+ CD3- CD11b- CD14- CD56- B220+ CD138- CD23+ CD21/35+ IgM+ |
| Marginal Zone | CD45+ CD19+ CD3- CD11b- CD14- CD56- B220+ CD93- CD23- CD21/35+ |
| Follicular | CD45+ CD19+ CD3- CD11b- CD14- CD56- B220+ CD93- CD23+ CD21/35+ |
| Germinal Center | CD45+ CD19+ CD3- CD11b- CD14- CD56- B220+ GL7+ CD95+ |
| Plasmablast | CD45+ CD19+ CD3- CD11b- CD14- CD56- B220+ CD138+ MHC II+ |
| Plasma cell | CD45+ CD19- CD3- CD11b- CD14- CD56- B220- CD138+ MHC II- |
| CD4+ T cell | CD45+ CD4+ TCRb+ |
| CD4+ Tfh | CD45+ CD4+ TCRb+ CXCR5+ PD1+ CD278(ICOS)+ |

| **Table S3: ADTs in Biolegend's Totalseq C Universal cocktail excluding isotype controls** | | | |
|---|---|---|---|
| S.No | ADT label | Common marker name | Gene (mouse) |
| 1 | HuMs.CD11b | CD11b (integrin αM) | Itgam |
| 2 | HuMs.CD44 | CD44 | Cd44 |
| 3 | HuMs.CD45R-B220 | CD45R/B220 (CD45 isoform) | Ptprc |
| 4 | HuMs.CD49f | CD49f (integrin α6) | Itga6 |
| 5 | HuMs.integrin.b7 | Integrin β7 | Itgb7 |
| 6 | HuMs.KLRG1 | KLRG1 | Klrg1 |
| 7 | HuMsRt.CD27 | CD27 | Cd27 |
| 8 | HuMsRt.CD278 | ICOS (CD278) | Icos |
| 9 | Ms.CCR3 | CCR3 | Ccr3 |
| 10 | Ms.CD103 | CD103 (integrin αE) | Itgae |
| 11 | Ms.CD106 | VCAM-1 (CD106) | Vcam1 |
| 12 | Ms.CD107a | LAMP-1 (CD107a) | Lamp1 |
| 13 | Ms.CD115 | CSF1R (CD115) | Csf1r |
| 14 | Ms.CD11a | CD11a (integrin αL) | Itgal |
| 15 | Ms.CD11c | CD11c (integrin αX) | Itgax |
| 16 | Ms.CD120b | TNFR2 (CD120b) | Tnfrsf1b |
| 17 | Ms.CD127 | IL-7Rα (CD127) | Il7r |
| 18 | Ms.CD134 | OX40 (CD134) | Tnfrsf4 |
| 19 | Ms.CD137 | 4-1BB (CD137) | Tnfrsf9 |
| 20 | Ms.CD138 | Syndecan-1 (CD138) | Sdc1 |
| 21 | Ms.CD150 | SLAMF1 (CD150) | Slamf1 |
| 22 | Ms.CD155 | PVR (CD155) | Pvr |
| 23 | Ms.CD160 | CD160 | Cd160 |
| 24 | Ms.CD163 | CD163 | Cd163 |
| 25 | Ms.CD169 | Siglec-1 (CD169) | Siglec1 |
| 26 | Ms.CD170 | Siglec-F (commonly labeled CD170 in mouse) | Siglecf |
| 27 | Ms.CD172a | SIRPα (CD172a) | Sirpa |
| 28 | Ms.CD182 | CXCR2 (CD182) | Cxcr2 |
| 29 | Ms.CD183 | CXCR3 (CD183) | Cxcr3 |
| 30 | Ms.CD185 | CXCR5 (CD185) | Cxcr5 |
| 31 | Ms.CD186 | CXCR6 (CD186) | Cxcr6 |
| 32 | Ms.CD19 | CD19 | Cd19 |
| 33 | Ms.CD1d | CD1d | Cd1d1/Cd1d2 |
| 34 | Ms.CD2 | CD2 | Cd2 |
| 35 | Ms.CD20 | CD20 | Ms4a1 |
| 36 | Ms.CD200 | CD200 | Cd200 |

| | | | |
|---|---|---|---|
| 37 | Ms.CD200R | CD200R | Cd200r1 |
| 38 | Ms.CD205 | DEC-205 (CD205) | Ly75 |
| 39 | Ms.CD21-CD35 | CR2/CR1 epitope (CD21/CD35) | Cr2 |
| 40 | Ms.CD22 | CD22 | Cd22 |
| 41 | Ms.CD223 | LAG-3 (CD223) | Lag3 |
| 42 | Ms.CD226-10E5 | DNAM-1 (CD226) | Dnam1 |
| 43 | Ms.CD23 | CD23 (FcεRII) | Fcer2a |
| 44 | Ms.CD24 | CD24 | Cd24a |
| 45 | Ms.CD25 | IL-2Rα (CD25) | Il2ra |
| 46 | Ms.CD267 | TACI (CD267) | Tnfrsf13b |
| 47 | Ms.CD272 | BTLA (CD272) | Btla |
| 48 | Ms.CD273 | PD-L2 (CD273) | Pdcd1lg2 |
| 49 | Ms.CD274 | PD-L1 (CD274) | Cd274 |
| 50 | Ms.CD279 | PD-1 (CD279) | Pdcd1 |
| 51 | Ms.CD3 | CD3ε (complex) | Cd3e |
| 52 | Ms.CD301b | CD301b | Mgl2 |
| 53 | Ms.CD304 | Neuropilin-1 (CD304) | Nrp1 |
| 54 | Ms.CD31 | PECAM-1 (CD31) | Pecam1 |
| 55 | Ms.CD317 | BST2 (Tetherin, CD317) | Bst2 |
| 56 | Ms.CD319 | SLAMF7 (CD319) | Slamf7 |
| 57 | Ms.CD357 | GITR (CD357) | Tnfrsf18 |
| 58 | Ms.CD366 | TIM-3 (CD366) | Havcr2 |
| 59 | Ms.CD371 | CLEC12A (MICL, CD371) | Clec12a |
| 60 | Ms.CD38 | CD38 | Cd38 |
| 61 | Ms.CD4 | CD4 | Cd4 |
| 62 | Ms.CD40 | CD40 | Tnfrsf5 |
| 63 | Ms.CD41 | Integrin αIIb (CD41) | Itga2b |
| 64 | Ms.CD43 | Sialophorin (CD43) | Spn |
| 65 | Ms.CD45 | CD45 | Ptprc |
| 66 | Ms.CD45.2 | CD45.2 (Ptprc allele) | Ptprc |
| 67 | Ms.CD45RB | CD45RB isoform | Ptprc |
| 68 | Ms.CD48 | SLAMF2 (CD48) | Slamf2 |
| 69 | Ms.CD49a | Integrin α1 (CD49a) | Itga1 |
| 70 | Ms.CD49b | Integrin α2 (CD49b/DX5) | Itga2 |
| 71 | Ms.CD49d | Integrin α4 (CD49d) | Itga4 |
| 72 | Ms.CD5 | CD5 | Cd5 |
| 73 | Ms.CD51 | Integrin αV (CD51) | Itgav |
| 74 | Ms.CD54 | ICAM-1 (CD54) | Icam1 |
| 75 | Ms.CD55 | DAF (CD55) | Cd55 |
| 76 | Ms.CD62L | L-selectin (CD62L) | Sell |

| | | | |
|---|---|---|---|
| 77 | Ms.CD63 | CD63 | Cd63 |
| 78 | Ms.CD69 | CD69 | Cd69 |
| 79 | Ms.CD71 | Transferrin receptor (CD71) | Tfrc |
| 80 | Ms.CD73 | 5′-Nucleotidase (CD73) | Nt5e |
| 81 | Ms.CD80 | B7-1 (CD80) | Cd80 |
| 82 | Ms.CD86 | B7-2 (CD86) | Cd86 |
| 83 | Ms.CD88 | C5aR1 (CD88) | C5ar1 |
| 84 | Ms.CD8a | CD8α | Cd8a |
| 85 | Ms.CD8b | CD8β | Cd8b1 |
| 86 | Ms.CD9 | CD9 | Cd9 |
| 87 | Ms.CD90.2 | Thy-1.2 (CD90.2) | Thy1 |
| 88 | Ms.CD93 | CD93 | Cd93 |
| 89 | Ms.CD94 | CD94 | Klrd1 |
| 90 | Ms.CD95 | Fas (CD95) | Fas |
| 91 | Ms.CD98 | 4F2hc (CD98) | Slc3a2 |
| 92 | Ms.F4-80 | F4/80 | Adgre1 |
| 93 | Ms.FceRIa | FcεRI α chain | Fcer1a |
| 94 | Ms.FR4 | Folate receptor 4 (FR4) | Folr4 |
| 95 | Ms.I.A-I.E | MHC class II I-A/I-E | H2-Aa/H2-Ab1/H2-Eb1 |
| 96 | Ms.IgD | Ig heavy chain δ | Ighd |
| 97 | Ms.IgG1 | Ig heavy chain γ1 | Ighg1 |
| 98 | Ms.IgG2b | Ig heavy chain γ2b | Ighg2b |
| 99 | Ms.IgM | Ig heavy chain μ | Ighm |
| 100 | Ms.Ly.49A | Ly49A | Klra1 |
| 101 | Ms.Ly.6A-E | Sca-1 (Ly6A/E epitope) | Ly6a (±Ly6e) |
| 102 | Ms.Ly.6C | Ly6C | Ly6c1/Ly6c2 |
| 103 | Ms.Ly.6G | Ly6G | Ly6g |
| 104 | Ms.Ly108 | Ly108 (SLAMF6) | Slamf6 |
| 105 | Ms.Ly49D | Ly49D | Klra4 |
| 106 | Ms.Ly49H | Ly49H | Klra8 |
| 107 | Ms.NK.1.1 | NK1.1 epitope | Klrb1c (strain-dependent) |
| 108 | Ms.Siglec-H | Siglec-H | Siglech |
| 109 | Ms.TCR.Bchain | TCR β chain (constant) | Trbc1/Trbc2 |
| 110 | Ms.TCR.GD-UC7 | TCR γδ | Trdc/Trgc (family) |
| 111 | Ms.TCR.RD-GL3 | TCR δ | Trdc |
| 112 | Ms.TCR.Va2 | TCR Vα2 | Trav2 family |
| 113 | Ms.TCR.Va8.3-KT50 | TCR Vα8.3 | Trav8-3 |
| 114 | Ms.TCR.Vb5.1-5.2 | TCR Vβ5.1/5.2 | Trbv5-1/Trbv5-2 |
| 115 | Ms.TCR.Vb8.1-8.2 | TCR Vβ8.1/8.2 | Trbv8-1/Trbv8-2 |

| | | | |
|---|---|---|---|
| 116 | Ms.TER.119 | TER-119 (Ly-76 antigen) | Ly76 |
| 117 | Ms.Tim.4 | TIM-4 | Timd4 |
| 118 | Ms.VISTA | VISTA | Vsir |
| 119 | MsRt.CD29 | CD29 (integrin β1) | Itgb1 |
| 120 | MsRt.CD61 | CD61 (integrin β3) | Itgb3 |
| 121 | MsRt.CD81 | CD81 | Cd81 |

**Table S4. Additional Spike-in CITEseq Antibodies**

| S. No | Marker | TotalSeq-C Spike-in Antibodies Cat# |
|---|---|---|
| 1 | CD273 (PD-L2) | 107229 |
| 2 | CD80 | 104755 |
| 3 | CD95 | 152616 |
| 4 | IgM | 406541 |
| 5 | CD267 | 133405 |
| 6 | CD319 (SLAMF7) | 152007 |
| 7 | IgG2b | 406720 |
| 8 | IgG1 | 406636 |

**Table S5. Mouse splenocyte sample pools for single cell CITE-seq analysis**

| **Sample** | **Mouse #** | **Mouse Hashtags** |
|---|---|---|
| Splenocytes 4540 | 34,35, 36, 37, 38 | Hash-tag1,5, 11, 7, 15 (Cat # 155861, 155869, 155881, 155873, 155889 respectively) |
| Splenocytes POV | 44, 46, 47, 48 | Hash-tag2, 12, 8, 16 (Cat # 155863, 155883, 155875, 155891 respectively) |
| Splenocytes ATA | 54, 56, 58 | Hash-tag3, 14, 1(Cat # 155865, 155887, 155861 respectively) |
| Splenocytes MOTA | 64, 65, 66, 67, 68 | Hash-tag4, 6, 13, 9, 2 (Cat # 155867, 155871, 155885, 155877, 155863 respectively) |
